# Intestinal stem cell marker MEX3A regulates PPARγ expression with functional impact in colorectal carcinogenesis

**DOI:** 10.1101/2025.01.13.632758

**Authors:** Ana R. Silva, Alexandre Coelho, Vanessa Machado, Morgana Russel, Dalila Mexieiro, Ana L. Amaral, Bruno Cavadas, Carina Carvalho-Maia, Davide Gigliano, Carmen Jerónimo, Raquel Almeida, Bruno Pereira

## Abstract

RNA-binding proteins (RBPs) are major effectors of post-transcriptional regulation. Recently, we described the role of MEX3A in maintaining intestinal stem cell identity and epithelial renewal by repressing the PPARγ pathway. This work aimed to study MEX3A functional impact in colorectal cancer (CRC). MEX3A and PPARγ expression profiles were characterized in murine and human models. CRISPR/Cas9-mediated MEX3A knockout was performed in patient-derived CRC tumoroids (PDCTs) and MEX3A RNA targets identified through the HyperTRIBE technique. *Apc^+/fl^*;*Mex3a^+/−^* mice presented a significant reduction in tumor burden. *Apc^+/fl^*;*Kras^+/G12D^*;*Mex3a^+/−^*mice presented a reduced tumor area, while corresponding tumoroids exhibited reduced growth and enhanced differentiation potential mediated by PPARγ signalling. MEX3A overexpression (85% of human CRC cases) was inversely correlated with PPARγ downregulation (72% of cases). Accordingly, MEX3A-depleted PDCTs showed decreased *LGR5* expression, accompanied by increased PPARγ expression and higher sensitivity to 5-Fluorouracil/Oxaliplatin (FOLFOX)-based chemotherapy. The HyperTRIBE results revealed a direct interaction between MEX3A and *PPARG* transcripts.

**STATEMENT OF SIGNIFICANCE:** These results emphasize that MEX3A plays a crucial role in colorectal carcinogenesis, partially through regulation of the PPARG pathway, mediating tumour development and response to therapy, thus constituting a potential therapeutic target.

## INTRODUCTION

RNA-binding proteins (RBPs) are the main mediators of the post-transcriptional regulatory machinery and are involved in virtually every RNA molecule-related step, including processing, localization, stability, translation, and turnover.^1, 2^ Together with non-coding RNAs they form ribonucleoprotein complexes that coordinate an additional and reversible layer of protein regulation.^3, 4^ RBPs recognize specific structural features and/or motifs in transcripts through well-defined RNA-binding domains, such as K homology (KH) domains.^5^ Disturbances of their fine-tune functions underlie several pathological conditions,^6^ and there is growing evidence that tumor cells highjack RBP- mediated processes with repercussions in cancer hallmarks.^7, 8^ Colorectal cancer (CRC) remains the third most frequent cancer worldwide and the second most prevalent cause of cancer-related deaths. Hence, the description of new biomarkers with prognostic value or amenable to targeted therapies is extremely important.

The Mex-3 RNA Binding Family Member A (MEX3A) belongs to the evolutionary-conserved MEX-3 family of RBPs.^9^ In vertebrates, the family has four homologous genes (*MEX3A* to *MEX3D*), encoding proteins that are 95% identical between human and mouse.^10^ These possess two KH domains that mediate RNA target binding and a really interesting new gene (RING) finger domain with E3 ubiquitin ligase activity.^11, 12^ MEX3A colocalizes with the bona-fide intestinal stem cell (ISC) marker Leucine-rich repeat-containing G-protein coupled receptor 5 (LGR5) during intestinal homeostasis.^13, 14^ Through the characterization of the first *Mex3a* knockout (KO) mouse model, this protein was established as a functionally relevant ISC marker.^15^ *Mex3a* KO mice present retarded growth and a postnatal lethality phenotype, with a striking loss of the *Lgr5+* ISC pool, accompanied by upregulation of the peroxisome proliferator-activated receptor gamma (PPARγ) differentiation pathway.^15^ MEX3A has been shown to be overexpressed in several cancer types, including breast,^16^ brain,^12^ and lung.^17^ Recently, it was proposed that drug-tolerant persister CRC cells, which mediate relapse after chemotherapy, also express MEX3A.^18^ However, the functional impact of MEX3A in the colorectal carcinogenic process has not yet been systematically addressed, nor its RNA targets.

Through in-depth characterization of CRC mouse models, human CRCs, and *ex vivo* tumoroid cultures, we provide evidence that MEX3A plays a relevant role in colorectal carcinogenesis. MEX3A was overexpressed in the early stages of the carcinogenic cascade and was detected in murine *Apc* mutant colonic adenomas and *Apc*;*Kras* mutant colon adenocarcinomas. Importantly, its downregulation led to a significantly reduced tumor burden in both mouse models. Mouse-derived tumoroids with decreased *Mex3a* expression exhibited morphological and molecular changes reminiscent of a more differentiated phenotype, which was replicated upon treatment with a PPARγ agonist. In agreement with this, MEX3A expression was substantially increased in a cohort of stage II colon cancer patients and was significantly correlated with low PPARγ levels. Additionally, patient-derived tumoroids with CRISPR/Cas9- mediated MEX3A depletion showed decreased *LGR5* mRNA levels, accompanied by increased PPARγ protein expression and higher sensitivity to chemotherapy. Finally, by employing HyperTRIBE (Targets of RNA-binding proteins Identified By Editing) methodology, we defined a list of putative MEX3A RNA targets, demonstrating a direct interaction between MEX3A and *PPARG* transcripts. Taken together, these findings revealed that the MEX3A/PPARγ regulatory axis is relevant for CRC development and therapeutic outcomes.

## RESULTS

### Increased *Mex3a* expression and loss of PPARγ are early molecular events in colorectal carcinogenesis

To evaluate the impact of MEX3A in colorectal carcinogenesis and explore a putative association with the most common signalling pathways involved in this process, we first characterized the *Mex3a* mRNA expression profile in mouse models with *Apc* inactivation, *Kras* activation, and combined mutations (n=5). Animals carrying the corresponding *loxP* alleles were crossed with *Fabpl^+/cre^* mice to generate deletions specifically in the distal small intestine and colonic epithelia.

Histological analysis of the colonic mucosa of wild-type adult mice (Supplementary Fig. S1) revealed the anticipated delimitation between the differentiated and proliferative compartments, with the former comprising the top two-thirds of the crypts, as evidenced by the presence of AB+ mature goblet cells, and the latter comprising the bottom third of the crypt, as indicated by KI67 staining. *Mex3a* transcripts were restricted to the bottom of the crypt and localized within the spatial domain of *Lgr5*+ cells (Supplementary Fig. S1). In *Apc^+/fl^*;*Fabpl^+/cre^* mice, stochastic loss of heterozygosity at the *Apc* locus led to spontaneous development of focal adenomas in the ileum and colon (Fig. 1A). These lesions were clearly identifiable by Wnt pathway overactivation, as shown by the increased expression of nuclear β-Catenin, accompanied by increased proliferative capacity (ectopic KI67+ cells) and decreased cellular differentiation, as indicated by the reduction in the number of AB+ goblet cells (Fig. 1B). A striking increase in the number of *Lgr5*+ cells was observed within these lesions, which appeared to be mainly accumulated at the base of the adenomatous glands, a phenomenon previously described in other mouse models and human tissues.^19, 20^ Concomitantly, we observed an increased number of *Mex3a* mRNA expressing cells, with a higher expression level (red spot intensity) per tumor cell when compared to the adjacent normal colonic tissue (Fig. 1B). Previously, we reported MEX3A-mediated regulation of PPARγ signalling critical for intestinal homeostasis.^15^ Here, we observed that PPARγ expression was strongly reduced in adenomas (Fig. 1B).

**Figure 1.**
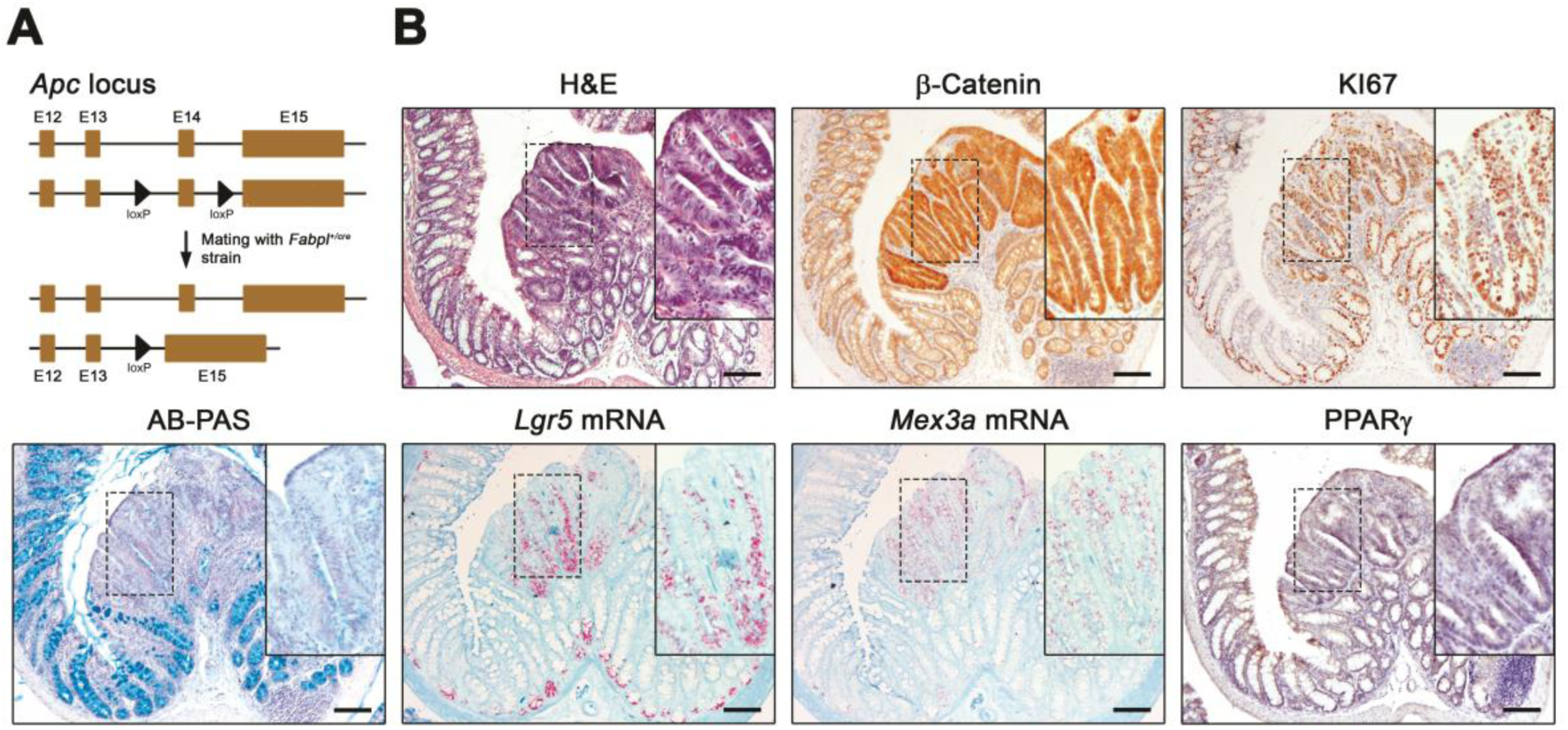
*Apc^+/fl^*;*Fabpl^+/cre^*mice colon adenomas show increased *Mex3a* levels and downregulated PPARγ expression. **A,** Representation of the genetically modified *Apc* locus for conditional inactivation by Cre recombinase. **B,** Histopathology of *Apc^+/fl^*;*Fabpl^+/cre^* adenomas (H&E and AB-PAS staining; β-Catenin, KI67 and PPARγ IHC; *Lgr5* and *Mex3a* mRNA ISH). Inserts depict high magnification of boxed areas. Scale bars, 100µm.

*Kras^+/G12D^*;*Fabpl^+/cre^* mice (Supplementary Fig. S2A) developed widespread hyperplasia throughout the colonic epithelium (Supplementary Fig. S2B), reminiscent of human hyperplastic polyps. This was characterized by crypt lengthening, mainly due to an extension of the KI67+ cell pool and an increased number of prominent AB+ goblet cells (Supplementary Fig. S2B). No differences in relation to the wild-type were observed concerning Wnt activity, *Lgr5*+ and *Mex3a*+ cells and PPARγ expression (Supplementary Fig. S2B).

When combined with *Apc* inactivation, *Kras* activation promotes the progression of adenomatous lesions to adenocarcinomas.^21^ Hence, colonic lesions in *Apc^+/fl^*;*Kras^+/G12D^*;*Fabpl^+/cre^*animals (Fig. 2A) were markedly different from those arising in *Apc* single mutants: they developed faster (animals were euthanized around 10 weeks of age compared to 30 weeks on average for *Apc* mutant mice) and were more extensive and heterogeneous (Fig. 2B). Strong accumulation of *Lgr5*+ and *Mex3a*+ cells was confined to tumor tissue showing high Wnt activity, as evidenced by increased β-Catenin nuclear expression, together with the presence of KI67+ cells and loss of AB+ cells (Fig. 2B). PPARγ expression was almost absent only in the areas with enhanced Wnt activity (Fig. 2B). Localized transformation events were also visible, which may indicate putative crypts of origin (Fig. 2C). These transformed foci, recognized by β-Catenin increased expression and loss of AB staining, showed *Mex3a*+ and *Lgr5*+ cells, with a concomitant decrease in PPARγ expression (Fig. 2C). The obtained results support the existence of a close molecular link between the initial genetic alteration most commonly observed in colorectal carcinogenesis (Wnt pathway augmented activity), increased *Mex3a* levels and decreased PPARγ expression.

**Figure 2.**
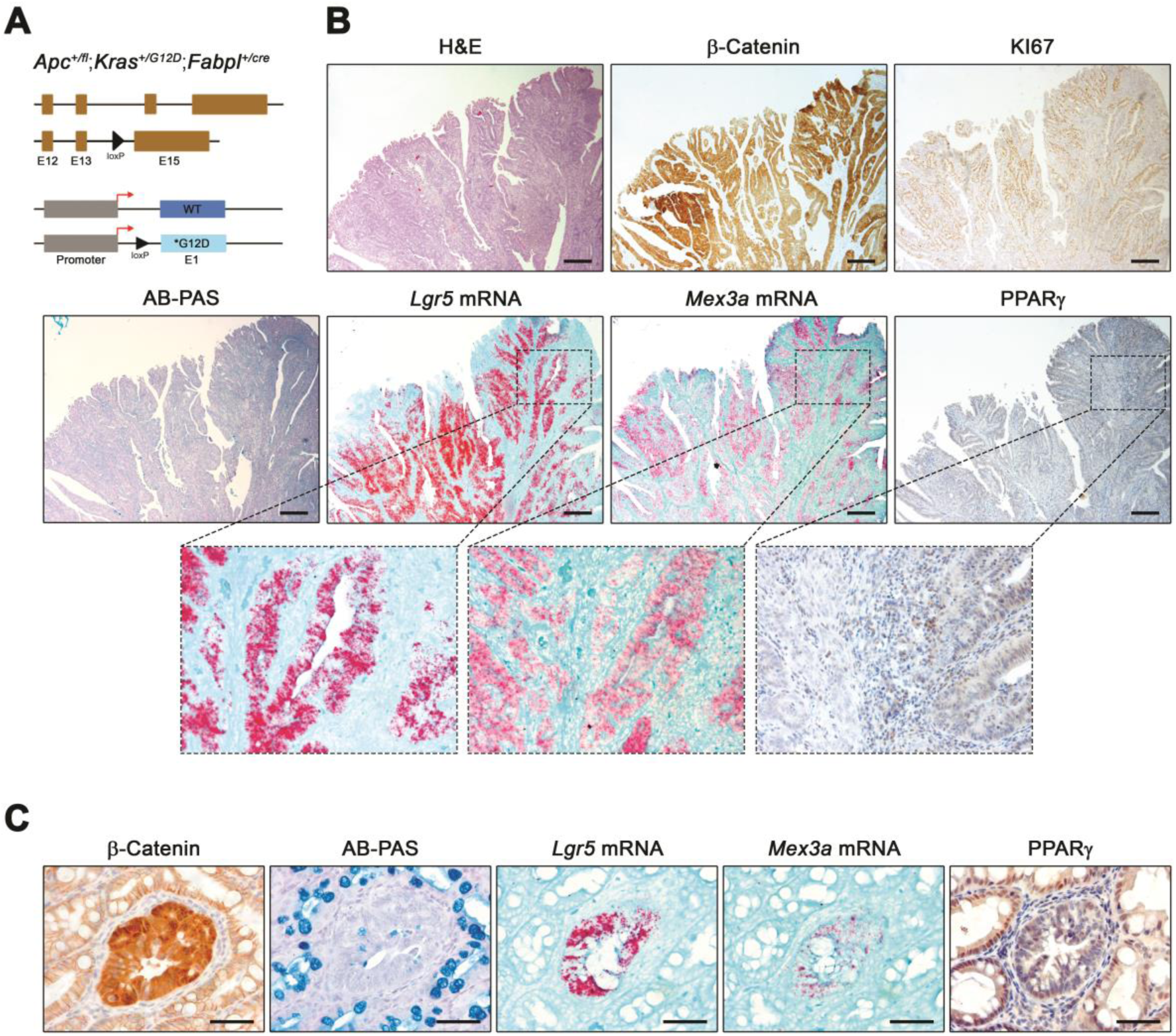
*Apc^+/fl^*;*Kras^+/G12D^*;*Fabpl^+/cre^*mice colon adenocarcinomas present strong accumulation of *Mex3a^+^*cells accompanied by loss of PPARγ expression. **A,** Representation of the modified *Apc* and *Kras* loci in the intestinal epithelium after crossing with *Fabpl^+/cre^* mice. **B,** Histopathology of *Apc^+/fl^*;*Kras^+/G12D^*;*Fabpl^+/cre^*adenocarcinomas (H&E and AB-PAS staining; β-Catenin, KI67 and PPARγ IHC; *Lgr5* and *Mex3a* mRNA ISH). Inserts depict high magnification of boxed areas. Scale bars, 200µm. **C,** Histopathology of single transformed foci in *Apc^+/fl^*;*Kras^+/G12D^*;*Fabpl^+/cre^*mice (AB-PAS staining; β-Catenin and PPARγ IHC; *Lgr5* and *Mex3a* mRNA ISH). Scale bars, 50µm.

### *Mex3a* heterozygosity impairs adenoma formation in an *Apc*-inactivation background

To address the role of MEX3A in adenoma initiation, independent cohorts of age-matched *Apc^+/fl^*;*Fabpl^+/cre^*;*Mex3a^+/−^* and *Apc^+/fl^*;*Fabpl^+/cre^* animals (n=15) were obtained. We observed that *Apc^+/fl^*;*Fabpl^+/^*^cre^ mice exhibited signs of disease more frequently than *Apc^+fl^*;*Fabpl^+/cre^*;*Mex3a^+/−^*littermates (60% versus 33%, respectively; Fig. 3A). The number of lesions in the small intestine and colon was quantified, revealing a significant decrease (*P* = 0.018) in the tumor burden in *Apc^+/fl^*;*Fabpl^+/cre^*;*Mex3a^+/−^*mice, specifically in the ileum (Fig. 3B). While 20% (3/15) of the *Apc^+/fl^*;*Fabpl^+/cre^*;*Mex3a^+/−^*animals had no lesions, and 67% (10/15) showed one to three tumors in the terminal part of the small intestine, all *Apc^+/fl^*;*Fabpl^+/cre^* mice developed lesions, with 67% (10/15) having four or more tumors. The developed lesions showed increased expression of nuclear β-Catenin, together with the presence of KI67+ cells and the absence of AB+ goblet cells (Fig. 3C). *Lgr5* and *Mex3a* expression levels were also increased compared to the adjacent normal-like tissue (Fig. 3C). Simultaneously, PPARγ expression was absent in the transformed cells (Fig. 3C). On the other hand, the tumor burden in the colon was similar for both genotypes, with about half of all animals not presenting lesions (Supplementary Fig. S3A and S3B). These data imply that MEX3A expression is not absolutely required for Wnt-mediated tumor initiation, but that its reduced expression and/or function impairs tumor growth.

**Figure 3.**
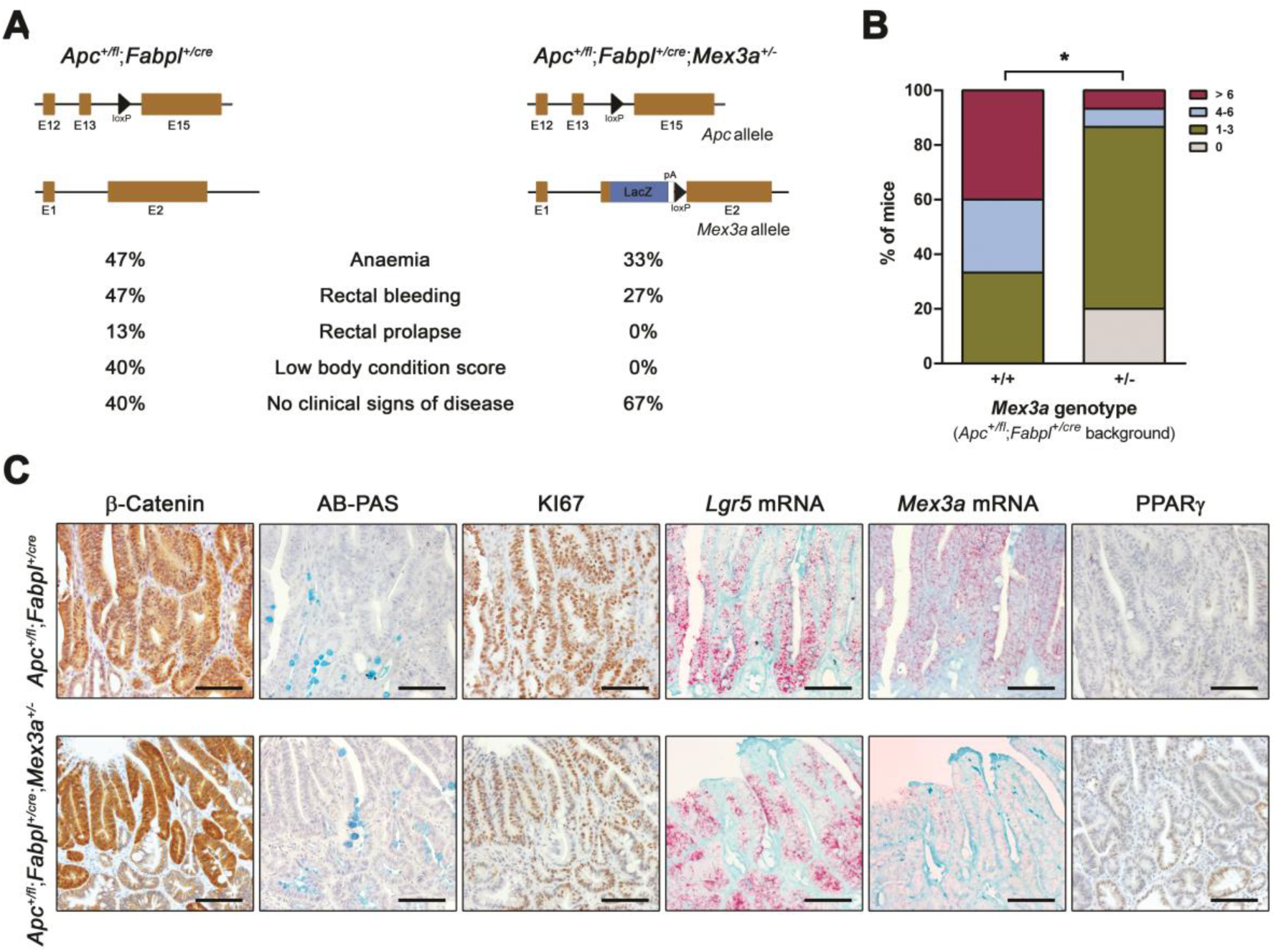
*Mex3a* downregulation impairs adenoma formation in *Apc^+/fl^*;*Fabpl^+/cre^* mice. **A,** Percentage of animals with different *Mex3a* genotypes in an *Apc^+/fl^*;*Fabpl^+/cre^* background presenting signs of disease (n=15). **B,** Quantification of the number of tumours in the ileum of *Apc^+/fl^*;*Fabpl^+/cre^*and *Apc^+/fl^*;*Fabpl^+/cre^*;*Mex3a^+/−^*mice. \**P* = 0.0182, Chi-square test. **C,** Histopathology of *Apc^+/fl^*;*Fabpl^+/cre^* and *Apc^+/fl^*;*Fabpl^+/cre^*;*Mex3a^+/−^* ileal lesions (AB-PAS staining; β-Catenin, KI67 and PPARγ IHC; *Lgr5* and *Mex3a* mRNA ISH). Scale bars, 100µm.

### *Mex3a* downregulation reduces CRC burden in an *Apc^+/fl^*;*Kras^+/G12D^*background and impacts CRC tumoroids growth dynamics

Due to the development of few colonic lesions in the *Apc* mutant strains, we established independent cohorts of age-matched *Apc^+/fl^*;*Kras^+/G12D^*;*Fabpl^+/cre^*;*Mex3a^+/−^*and *Apc^+/fl^*;*Kras^+/G12D^;Fabpl^+/cre^*animals (n=7 and n=4, respectively). Macroscopic examination revealed a 43% decrease (*P* = 0.0021) in tumor growth area in the distal colon of *Apc^+/fl^*;*Kras^+/G12D^*;*Fabpl^+/cre^*;*Mex3a^+/−^*mice (0.99±0.17cm^2^) when compared to *Apc^+/fl^*;*Kras^+/G12D^;Fabpl^+/cre^*control animals (1.73±0.42cm^2^, Fig. 4A and B). Lesions were not observed in the small intestine.

**Figure 4.**
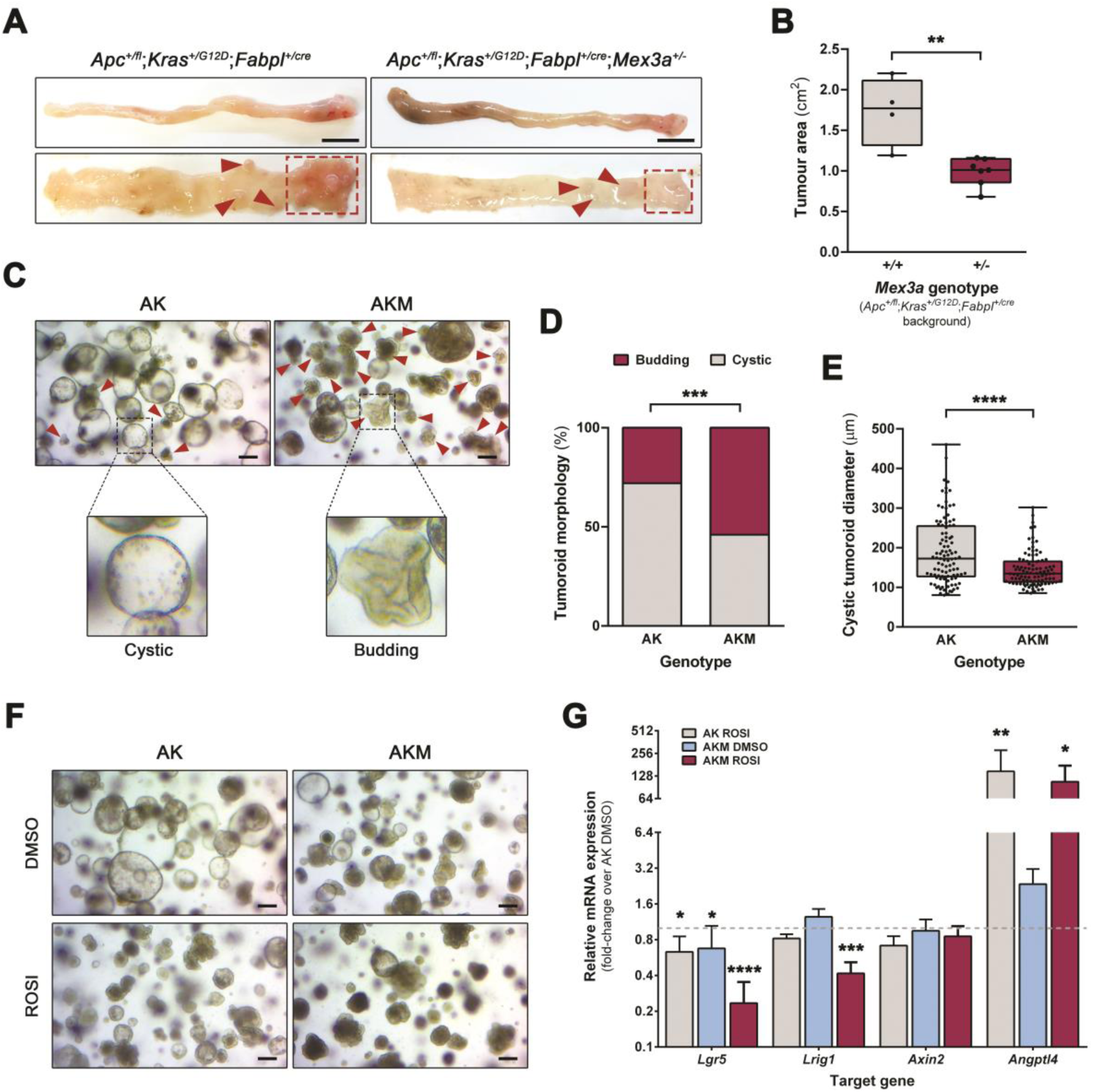
*Apc^+/fl^*;*Kras^+/G12D^*;*Fabpl^+/cre^;Mex3a^+/−^* mice present lower tumour burden and altered tumoroid growth dynamics. **A,** Macroscopic images of the colon from *Apc^+/fl^*;*Kras^+/G12D^*;*Fabpl^+/cre^*and *Apc^+/fl^*;*Kras^+/G12D^*;*Fabpl^+/cre^;Mex3a^+/−^*mice. Boxed areas and red arrows indicate tumours location. Scale bars, 1cm. **B,** Quantification of tumour area in *Apc^+/fl^*;*Kras^+/G12D^*;*Fabpl^+/cre^*and *Apc^+/fl^*;*Kras^+/G12D^*;*Fabpl^+/cre^;Mex3a^+/−^*colon (n=4 and n=7, respectively). Data are represented in a box-and-whisker plot as mean (middle line) with the minimum and maximum distribution values. Each point depicts one tumour. \*\**P* = 0.0021, unpaired Student’s *t*-test. **C,** Phase-contrast microscopy images of AK and AKM colon tumoroids after 7 days of culture. Red arrows indicate budding structures and inserts depict high magnification of boxed regions. Scale bars, 100µm. **D,** Quantification of the proportion of budding and cystic morphologies in AK and AKM tumoroids (n=3). \*\*\**P* = 0.0003, Fisher’s exact test. **E,** Quantification of the size (diameter length) of AK and AKM cystic tumoroids (n=3). Data are represented in a box-and-whisker plot as mean (middle line) with the minimum and maximum distribution values. Each point depicts one tumoroid. \*\*\*\**P* < 0.0001, Mann-Whitney test. **F,** Phase-contrast microscopy images of AK and AKM cultures after 5 days of treatment with 50µM ROSI or equal volume of the drug vehicle (DMSO). Scale bars, 100µm. **G,** Reverse transcription quantitative PCR analysis of *Lgr5*, *Lrig1*, *Axin2* and *Angptl4* mRNA expression in AK and AKM tumoroid cultures after ROSI treatment (n=3). Data is presented as the mean fold- change plus standard deviation for target gene expression relative to AK DMSO levels (dashed line). *Lgr5*: AK ROSI \**P* = 0.0136, AKM DMSO \**P* = 0.0317, AKM ROSI \*\*\*\**P* < 0.0001; *Lrig1*: AKM ROSI \*\*\**P* = 0.008; *Angptl4*: AK ROSI \*\**P* = 0.0037, AKM ROSI \**P* = 0.0376, all one-way ANOVA.

To directly examine MEX3A role in cancer cell behaviour, we established *Apc^+/fl^*;*Kras^+/G12D^*;*Fabpl^+/cre^*;*Mex3a^+/−^*(AKM)- and *Apc^+/fl^*;*Kras^+/G12D^*;*Fabpl^+/cre^*(AK)-derived tumoroid cultures (n=3). These were kept in culture media solely containing the growth factor Noggin for the selective expansion of tumor cells. AKM cultures had a significantly higher proportion of budding tumoroids (*P* = 0.0003), in contrast to the predominance of cystic structures in AK cultures (Fig. 4C, red arrows and inserts). More than half of the AKM tumoroids were morphologically reminiscent of normal budding organoids (Fig. 4D) suggesting increased differentiation. When comparing the cyst-like structures only, AKM tumoroids were also significantly smaller (*P* < 0.0001) than the AK controls (Fig. 4E).

Given our previous observations concerning the *Mex3a* KO mouse model,^15^ we hypothesized that altered PPARγ signalling could underlie the tumoroid phenotypic changes. To address this, we treated tumoroid cultures with the PPARγ-selective agonist ROSI for 5 days. ROSI treatment induced an overall decrease in tumoroid size alongside a striking change in tumoroid morphology, namely, a reduction in the number of cystic structures, mostly noticeable in the AK cultures (Fig. 4F). Treatment efficiency was confirmed by increased *Angptl4* mRNA expression levels, a PPARγ transcriptional target, whose expression was already endogenously enhanced in AKM cultures (Fig. 4G). Interestingly, ROSI treatment induced a marked decrease in the expression levels of *Lgr5* and *Lrig1* ISC markers, particularly in AKM tumoroids (*P* < 0.0001 and *P* = 0.0008, respectively; Fig. 4G). This effect was not due to Wnt signalling blockade, since the expression of the canonical Wnt target gene *Axin2* did not change (Fig. 4G). These results show that *Mex3a* downregulation decreases CRC growth *in vivo* and *ex vivo*, possibly by inducing a more differentiated phenotype through increased PPARγ activity.

### MEX3A is overexpressed in human CRC tissues

To explore the MEX3A expression profile in human CRC cases, we performed bioinformatics analysis using data from the cancer data-mining platform TCGA and the non-disease state GTEx portal. This revealed significantly increased *MEX3A* mRNA expression levels (*P* < 0.01) in both colon (COAD) and rectal (READ) adenocarcinomas compared to normal tissues (Supplementary Fig. S4A). The same significant difference was observed for *LGR5* (*P* < 0.01; Supplementary Fig. S4B). To obtain insights into MEX3A expression and tissue distribution in early cancer stages, we analyzed MEX3A protein levels by IHC in a cohort of stage II colon cancer patients (n=172). In the normal colon tissue (n=5), MEX3A staining was faintly detected in the lower crypt regions (Fig. 5A). In contrast, in 84.9% (146/172) of tumor tissues, MEX3A showed increased expression, with moderate to strong staining in over 66% of the cancer cells, mainly at the nuclear level (Fig. 5B). In the remaining 15.1% (26/172) of cases, MEX3A showed a weak staining pattern, with few negative cases (Fig. 5B). High MEX3A expression was significantly associated with Microsatellite Stable (MSS) tumors (*P* = 0.007; Supplementary Table 1). No significant associations were found with other clinicopathological parameters.

**Figure 5.**
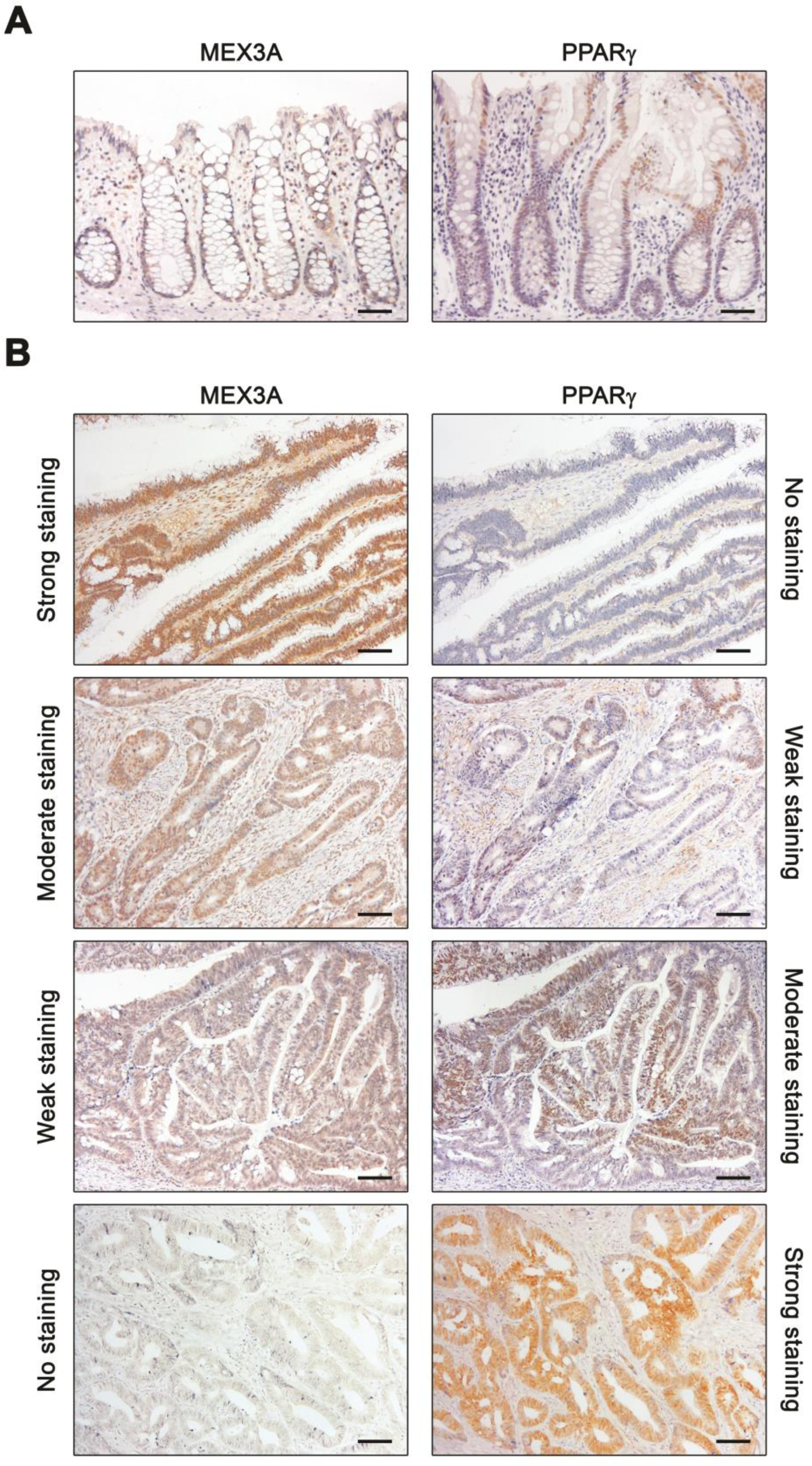
MEX3A and PPARγ are differentially expressed in human colon tumours. **A,** IHC for MEX3A and PPARγ in human normal colon tissues. Scale bars, 50µm. **B,** IHC for MEX3A and PPARγ in cancer cases, portraying tumours with strong, moderate, weak and negative staining. Scale bars, 50µm.

Additionally, we analyzed PPARγ levels in the same series. In normal colonic tissues, PPARγ was detected mainly in the upper crypt regions (Fig. 5A). In contrast to MEX3A, 72.1% (124/172) of the cases were negative or weak for PPARγ expression, whereas 26.7% (46/172) presented moderate to strong staining (Fig. 5B). High PPARγ levels were also significantly associated with MSS tumors (*P* < 0.0001), but no statistically significant associations were observed with other parameters (Supplementary Table 1). Despite the observed correlations with MSS status and possible implications for patient response to treatment, MEX3A and PPARγ expression profiles did not correlate with patient outcome (Supplementary Fig. S4C and S4D). Interestingly, we were able to establish a statistically significant correlation between high MEX3A levels and lower PPARγ expression (*P* = 0.039). In line with our findings in murine CRC models, the data sustains that there is also an association between increased MEX3A expression and downregulated PPARγ in the early stages of human colorectal carcinogenesis.

### *MEX3A* KO in patient-derived CRC tumoroids leads to re-expression of PPARγ and enhances sensitivity to chemotherapy

To gather functional evidence of the role of MEX3A in human colon cancer cells, we established patient-derived CRC tumoroid (PDCT) lines (Supplementary Fig. S5A). In agreement with the CRC cases previously analyzed, we observed an inverse correlation between MEX3A and PPARγ protein expression profiles in both PDCTs and normal colon-derived organoid lines (Fig. 6A and B and Supplementary Fig. S5B). The PDCT line 005T was genetically engineered using CRISPR/Cas9 technology to establish isogenic human tumoroids with MEX3A KO (005T_AE6; Fig. 6C and D, and Supplementary Fig. S5C). The lack of MEX3A did not seem to significantly alter tumoroid growth kinetics compared to the parental line (Fig. 6E and F). However, at late culture stages (around 2 weeks), MEX3A KO PDCTs had thicker walls (Fig. 6G), became darker in appearance due to accumulation of debris inside the lumen and underwent cell death, mostly those positioned deeper inside the Matrigel dome (Fig. 6E). These features suggested an altered cellular differentiation profile.^22^ Most interestingly, MEX3A KO tumoroids exhibited a significant decrease in *LGR5* mRNA levels (*P* = 0.001; Fig. 6H and I), accompanied by a significant increase in PPARγ protein expression (Fig. 6D and H).

**Figure 6.**
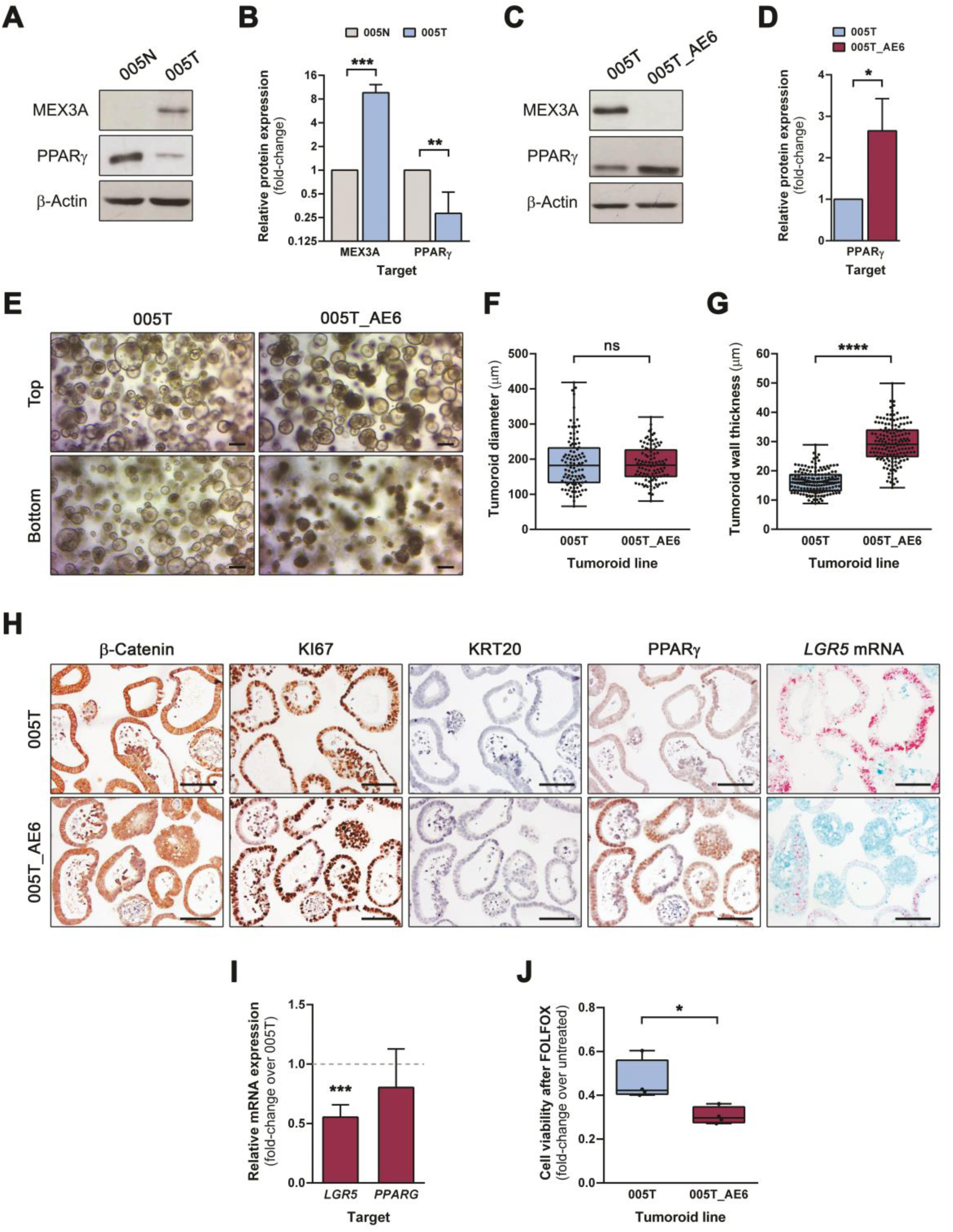
MEX3A KO in patient-derived CRC tumoroids increases PPARγ expression and enhances chemosensitivity. **A,** Western blot of MEX3A and PPARγ protein expression in 005T (tumoroids) and 005N (organoids) lines derived from the same patient. **B,** Quantification of MEX3A and PPARγ protein expression in 005T relative to 005N (n=4). Data is presented as mean fold-change plus standard deviation. \*\**P* = 0.001, \*\*\**P* = 0.0005, unpaired Student’s *t*-test. **C,** Western blot of MEX3A and PPARγ protein expression levels in 005T_AE6 (MEX3A KO) and parental 005T tumoroid lines. **D,** Quantification of PPARγ protein expression levels in 005T_AE6 relative to 005T (n=3). Data is presented as mean fold-change plus standard deviation. \**P* = 0.0211, unpaired Student’s *t*-test. **E,** Phase-contrast microscopy images of 005T and 005T_AE6 tumoroids at day 7 of culture, either near the top or the bottom of the Matrigel dome. Scale bars, 200µm. **F** and **G**, Quantifications of the size **(F)** and wall thickness **(G)** of 005T_AE6 and 005T cystic tumoroids (n=3). Data are represented in box-and-whisker plots as mean (middle line) with the minimum and maximum distribution values. Each point depicts one tumoroid. ns (not significant), \*\*\*\**P* < 0.0001, Mann-Whitney test. **H,** Histopathology of 005T_AE6 and 005T tumoroid lines (β-Catenin, KI67, KRT20 and PPARγ IHC; *LGR5* mRNA ISH). Scale bars, 100µm. **I,** Reverse transcription quantitative PCR analysis of *LGR5* and *PPARG* mRNA expression levels in 005T_AE6 and 005T tumoroid cultures after 14 days of culture (n=6). Data is presented as the mean fold-change plus standard deviation for target gene expression relative to 005T levels (dashed line). \*\*\**P* = 0.0001, unpaired Student’s *t*-test. **J,** Quantification of cell viability of 005T_AE6 and 005T tumoroid lines after FOLFOX-based treatment for 72h (n=4). Data are represented in a box-and-whisker plot as mean (middle line) with the minimum and maximum distribution values. \**P* = 0.0286, Mann-Whitney test.

As the characterization of MEX3A KO PDCTs pointed to lower stemness potential, we hypothesized that abrogation of MEX3A expression could affect the response of CRC cells to chemotherapy. Thus, we exposed MEX3A KO tumoroids and corresponding isogenic controls to a combination of 5-FU and Oxaliplatin (FOLFOX regimen) for 72h. We observed that MEX3A KO tumoroids showed significantly higher sensitivity to treatment (P = 0.0286), exhibiting lower cell viability than the parental line (Fig. 6J). Collectively, these results demonstrate that MEX3A is necessary for LGR5+ stem cell subpopulation maintenance in human CRC cells, with a functional impact on chemotherapy response.

### *PPARG* mRNA is a MEX3A target in CRC cells

In parallel with previous functional characterizations, we sought to obtain mechanistic insights into MEX3A regulation. To this end, we employed a cutting-edge technique called HyperTRIBE, based on the establishment of a fusion protein between the human MEX3A coding sequence and the hyperactive version of the catalytic domain (cd) of the *Drosophila* RNA-editing enzyme Adenosine Deaminase Acting on RNA (ADAR).^23^ Once expressed, it creates irreversible adenosine-to-inosine (A-to-I) RNA-editing events that are identified as adenosine-to-guanosine (A-to-G) mismatches near MEX3A-binding sites (Fig. 7A). Editing specificity is guaranteed by the lack of ADARcd RNA-binding capacity. To control for background editing, we generated a MEX3A RNA-binding mutant (MEX3AΔKH) by introducing mutations in the KH domain hallmark GxxG loops. MEX3A-ADARcd and MEX3AΔKH-ADARcd sequences were placed under the control of a doxycycline (DOX)-inducible promoter together with a TurboRFP reporter (Supplementary Fig. S6A). Upon lentiviral transduction of the human CRC cell line SW480 and establishment of puromycin-selected cell populations, we confirmed tight regulation of the system 48h after DOX addition by RFP fluorescence detection (Supplementary Fig. S6B) and MEX3A-ADARcd fusion protein overexpression (Supplementary Fig. S6C). RFP+ cells were collected (Supplementary Fig. S6D) and used for RNA-sequencing with the corresponding non-induced controls (NIC).

**Figure 7.**
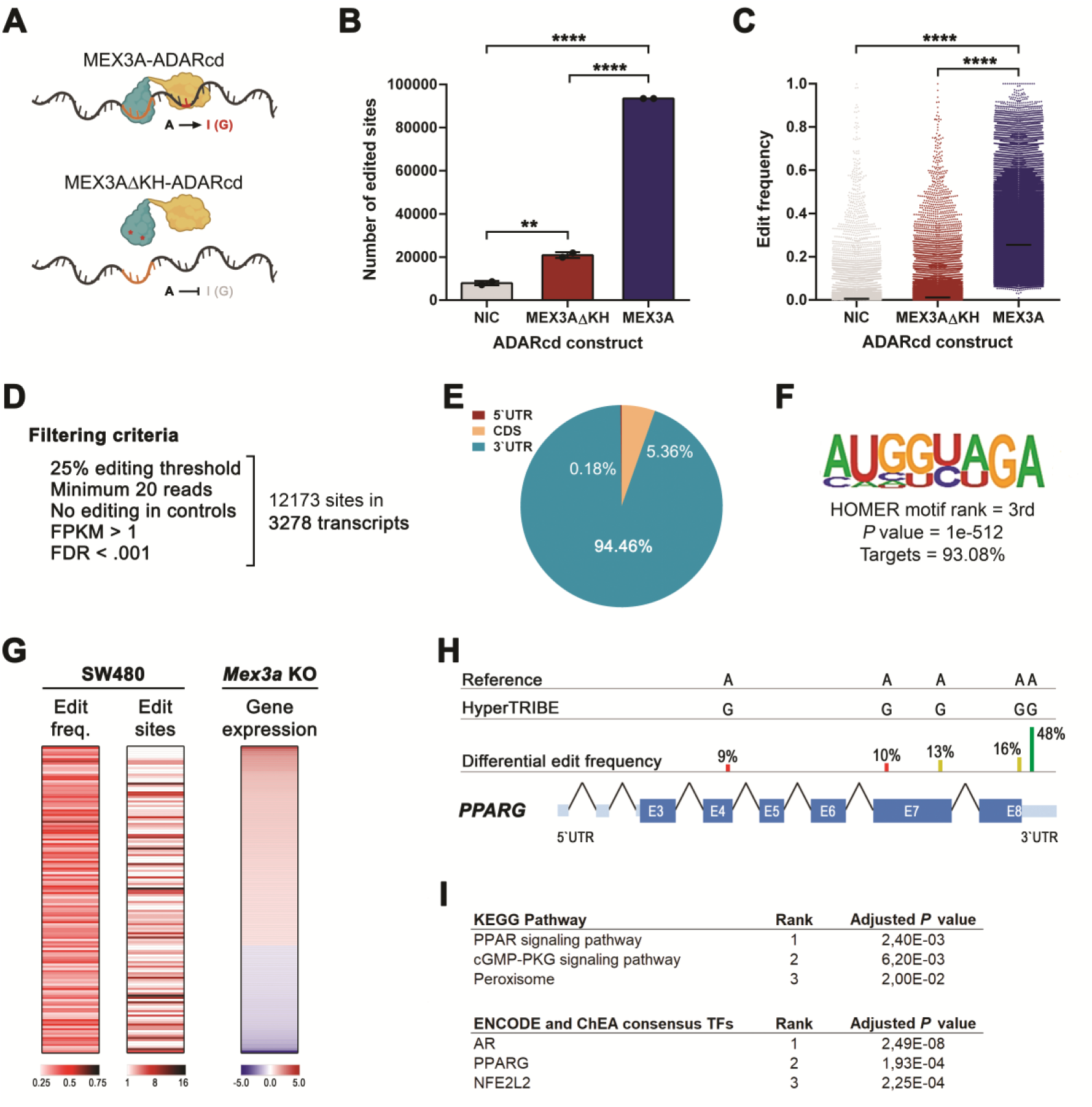
HyperTRIBE identifies *PPARG* as a MEX3A direct target in CRC cells. **A,** Representation of HyperTRIBE methodology, showing the MEX3A protein fused to the catalytical domain of ADAR (MEX3A-ADARcd, top) and the mutant control (MEX3AΔKH- ADARcd, bottom). **B,** Number of edit sites on mRNA transcripts. Data are represented as means plus standard deviation in two independent experiments. \*\**P* = 0.0019, \*\*\*\**P* < 0.0001, Tukey’s multiple comparisons test. **C,** Edit frequency distribution of the significant sites (*P* < 0.05). Each dot represents an edit event, with the mean value represented as a black line. \*\*\*\**P* < 0.0001, Dunn’s multiple comparisons test. **D,** Filtering criteria applied to the dataset. **E,** Distribution of the edit sites on the transcripts. **F,** *De novo* motif analysis. **G,** Heatmap of the 137 transcripts showing edit sites in SW480 cells and with significantly altered expression in *Mex3a* KO intestinal crypts. **H,** Distribution of the edit sites across the *PPARG* transcript. I, KEGG pathway and transcription factor analyses of the subset of 137 overlapping transcripts. The 3 most significant terms in each analysis are depicted according to their *P* value.

MEX3A-ADARcd protein induction in SW480 cells led to a significant increase in the absolute number of edited sites and the edit frequency of RNAs when compared to NIC or overexpression of the MEX3AΔKH-ADARcd mutant (Fig. 7B and C). Linear correlation analysis indicated that edit frequency was highly reproducible between replicates (Supplementary Fig. S6E) and independent of transcript size or transcript expression level (Supplementary Fig. S6F). Transient induction of the MEX3A-ADARcd fusion protein did not cause extensive transcriptional changes, with 90 transcripts being significantly altered (*P* < 0.01 and −2<fold-change>2, Supplementary Fig. S6G).

To confidently pinpoint MEX3A RNA targets, we applied a set of highly stringent filtering criteria that led to the identification of 12173 unique edited sites in 3278 transcripts (Fig. 7D). The majority of these sites (∼94%) were located in the 3’untranslated region (UTR), consistent with previous findings (Fig. 7E).^24, 25^ In addition, a *de novo* motif search using a 1000bp window centred at the edited sites revealed the presence of a particular sequence overlapping a known MEX-3 Recognition Element (MRE) within the top three most significantly enriched motifs (Fig. 7F).^24, 25^ We found that 137 edited transcripts were common to the *Mex3a* KO intestinal crypt gene signature,^15^ implying a direct regulatory effect (Fig. 7G). Strikingly, *PPARG* mRNA was part of the subset of edited RNAs also upregulated upon *Mex3a* deletion, exhibiting a major edited site in the 3’UTR (Fig. 7H). KEGG pathway and transcription factor analyses of this subset revealed significant enrichment for the PPAR pathway and PPARγ transcriptional targets, respectively (Fig. 7I). Overall, the data demonstrates that MEX3A-ADARcd protein editing and regulatory functions are fully dependent on MEX3A RNA-binding ability and that *PPARG* is a direct MEX3A target.

## DISCUSSION

RBPs are crucial for controlling homeostatic cellular processes and their altered functions are associated with cancer initiation and progression. Using CRC mouse models, human tumor tissues, and *ex vivo* tumoroid cultures, we provide evidence that MEX3A plays a biologically relevant role in CRC, particularly in the early stages, controlling the balance between LGR5+ cancer stem cell subpopulations and PPARγ-mediated cell differentiation, with an impact on tumor development and therapeutic response.

Characterization of *Apc* mutant mice revealed a close connection between the first event of malignant transformation and increased *Mex3a* levels. While *LGR5* is a target of the Wnt/β-Catenin pathway,^26^ there are currently no data supporting *MEX3A* being directly regulated by Wnt signalling. As such, the ectopic *Mex3a* upregulation observed in *Apc* mutant-derived adenomas and *Apc*;*Kras* mutant-derived adenocarcinomas presumably reflects a higher percentage of cellular subpopulations with stem cell-like features and not merely amplified β-Catenin transcriptional activation. In contrast, *Mex3a* haploinsufficiency significantly decreased the CRC burden in these mouse models, both in terms of the number and extent of the lesions. Similar tumor attenuation effects were recently reported in inflammation-induced CRC mouse models with either a constitutive^27^ or conditional^18^ *Mex3a* KO, indicating a common phenotypic outcome independent of tumorigenic insult.

The characterization of a cohort of patients with stage II colon cancer demonstrated an inverse correlation between MEX3A and PPARγ expression levels at early CRC stages. We were also able to establish a statistically significant association between high MEX3A levels and MSS status, which might indicate a prognostic value for MEX3A, since stage II CRC patients with MSS tumors have a worse prognosis when compared to highly immunogenic microsatellite instability (MSI) tumors.^28, 29^ Although we could not establish a correlation between MEX3A levels and patient survival, probably due to the low number of relapses in our cohort, several studies have been published in the last couple of years linking MEX3A overexpression with worse prognosis in multiple cancer types, including brain,^12, 30^ breast,^16, 31^ lung,^17^ liver,^32^ pancreatic,^33^ kidney,^34^ and colon cancers.^27^

We observed that MEX3A deficiency led to differences in *ex vivo* tumoroid growth and morphology patterns, as well as in the PPARγ expression profile. The alterations in AKM mouse-derived tumoroids and in MEX3A KO PDCTs were not fully identical, most probably due to their distinct mutational backgrounds. However, the appearance of multiple organoid differentiation features,^22, 35^ including frequent budding events, darker aspect with accumulation of apoptotic cells inside the lumen, and thick-walled cysts, all point to common changes in cancer cells, indicative of a more differentiated state. In agreement with this, we detected low *LGR5* mRNA levels upon partial or complete MEX3A loss. Notably, this is accompanied by increased PPARγ expression, and the previous phenotypic and molecular features are replicated upon treatment with a PPARγ-specific synthetic agonist, supporting a regulatory effect.

PPARγ is a member of the nuclear hormone receptor superfamily, identified as a master transcriptional determinant of adipogenesis and lipid metabolism.^36^ It has since been demonstrated to regulate cellular differentiation in various biological settings, including suppression of osteoblast differentiation from mesenchymal stem cells,^37^ promotion of osteoclast differentiation from hematopoietic stem cells,^38^ differentiation of bladder epithelial cells,^39^ as well as macrophage differentiation from monocytes.^40^ In the gut, PPARγ is primarily localized in the differentiated epithelial compartment (Figs. 1B and 5A).^41^ On the other hand, altered PPARγ activity has been causally associated with disease, including neurological disorders, chronic inflammation, and cancer.^42^ Several *in vitro* studies have demonstrated that PPARγ activation causes cell cycle arrest and differentiation of cancer cell lines, including those derived from lung,^43^ prostate,^44^ breast,^45^ pancreas^46^ and colon.^47^ Available *in vivo* data also supports a role for PPARγ as a tumor suppressor, with *Pparg* intestine-specific KO animals showing increased tumor incidence in the context of transgenic, chemically-mediated or inflammation-induced CRC.^48–50^ Moreover, a large cohort study suggests that PPARγ expression is independently associated with good prognosis in CRC.^51^ A complex and reciprocally opposing crosstalk between the canonical Wnt pathway and PPARγ signalling has been reported in different biological contexts. However, in our CRC models, MEX3A abrogation led to increased PPARγ expression even in a background of constitutive activation of the Wnt/β-Catenin pathway. This suggests that fine-tuning of PPARγ activity is particularly dependent on MEX3A and is at least partially uncoupled from Wnt signalling status.

We previously observed that overactivation of the PPARγ pathway dramatically impairs normal intestinal organoid growth and expansion due to the loss of *Lgr5*+ intestinal stem cells, particularly in MEX3A absence.^15^ In the CRC context, despite a significant reduction in *LGR5* levels, both ROSI-treated AKM tumoroids and MEX3A KO PDCTs retained their self-renewal ability (data not shown). This suggests that LGR5- cancer cell subpopulations with stem-like properties may contribute to tumor propagation upon MEX3A depletion. Nevertheless, complete ablation of MEX3A expression in human PDCTs instigates a heightened chemotherapeutic response to FOLFOX. In agreement, an association between partial MEX3A inhibition and improved chemotherapy results has been reported, namely for glioblastoma cells treated with temozolomide,^30^ pancreatic ductal adenocarcinoma cell lines upon gemcitabine treatment,^33^ hepatocellular carcinoma cell lines exposed to sorafenib,^32^ and also for a CRC patient-derived subcutaneous xenograft subjected to a combination of 5-FU and SN-38 (FOLFIRI regimen),^18^ thus strengthening the concept that MEX3A downregulation generates a tumor cell vulnerability that can be therapeutically exploited.

High-throughput identification of putative MEX3A targets was achieved using HyperTRIBE, providing the first MEX3A RNA interactome in CRC cells. To rule out potential off-target editing, we employed a MEX3A protein with impaired RNA-binding capacity. The MEX3AΔKH-ADARcd mutant produced a much lower number of edited sites and had a very low edit frequency, equivalent to the NIC. Most importantly, transcripts with any editing events detected in the MEX3AΔKH-ADARcd or NIC conditions were excluded from the target list. Successful adaptation of the technique was further supported by the specific enrichment of MRE-like motifs in the vicinity of edited sites in 93% of targets. To prevent analytical bias by the ADAR itself, we implemented a TET-On system and performed transient inductions. Differential gene expression analysis showed that MEX3AΔKH-ADARcd protein did not disturb the cell transcriptome (Supplementary Fig. S6G). Hence, the alterations observed upon MEX3A-ADARcd activation were exclusively due to MEX3A function. Interestingly, very few of the edited targets had altered expression levels in our experimental setup (only 73, considering *P* < 0.05 and −1.5<fold-change>1.5). This might be a consequence of short-term induction or may indicate a major role for MEX3A in translational control. Nevertheless, the results indicate that editing activity reflects MEX3A-specific binding.

In line with our *in vivo* and *ex vivo* data, *PPARG* mRNA is part of the list of MEX3A HyperTRIBE-edited transcripts. Although only one *PPARG* edited site passed the strict filtering criteria established, lowering the edit frequency to a still conservative 10% threshold^23^ revealed two additional high-confidence sites in the coding sequence, with 13% and 16% edit frequencies (Fig. 7H), strengthening *PPARG* transcript as a true MEX3A binding target. Other elements of the PPAR pathway also had editing events, including glycerol kinase (*GK*), sorbin and SH3 domain containing 1 (*SORBS1*), acyl-CoA oxidase 1 (*ACOX1*), and stearoyl-CoA desaturase 5 (*SCD5*). This is reminiscent of the post-transcriptional regulon model,^52^ whereby MEX3A may control the expression of functionally related transcripts in a coordinated manner to achieve fine-tuning of a specific biochemical process. Our future studies will address MEX3A effect over additional signalling pathways, highly represented within the set of MEX3A-ADARcd edited transcripts, and how the MEX3A-mediated regulatory network further contributes to colorectal carcinogenesis.

## MATERIALS AND METHODS

### Animal models

Animal experimentation was performed in accordance with the Portuguese National Regulation established by Decreto-Lei 113/2013 that is the national transposition of the European Directive 2010/63/EU for the Care and Use of Laboratory Animals. Mice were bred at the i3S Animal Facility, accredited by the Association for Assessment and Accreditation of Laboratory Animal Care (AAALAC). All strains were in a C57BL/6 background. The *Mex3a* KO strain was generated at the National Centre of Biotechnology (CNB-CSIC, Madrid, Spain) and previously characterized.^15^ The *Apc^2Lox14^*, *Kras^LSL-G12D^*, and *Fabpl^cre^*strains (a kind gift from Sérgia Velho, i3S, Portugal) were also previously established and characterized.^21^ The *Apc^2Lox14^*, *Kras^LSL-G12D^* and *Fabpl^cre^*strains were mated with the *Mex3a* heterozygous strain to obtain *Apc^+/fl^;Fabpl^+/cre^;Mex3a^+/−^*and *Apc^+/fl^*;*Kras^+/G12D^*;*Fabpl^+/cre^;Mex3a^+/−^*cohorts and corresponding *Apc^+/fl^;Fabpl^+/cre^* and *Apc^+/fl^*;*Kras^+/G12D^*;*Fabpl^+/cre^*controls. Animals were genotyped using specific primer pairs. Mixed cohorts were kept under a standard 12h light/dark cycle, with water and rodent chow available *ad libitum* and housed in Type II polycarbonate cages in single sex groups of two to six animals. Each cage was provided with corncob bedding (LBS serving Biotechnology, UK), sheets of absorbent paper, a cardboard tube (LBS serving Biotechnology, UK) for nesting and an acrylic tunnel for handling. To determine the number of animals to be used for tumour development evaluation in different genetic backgrounds we used the G*Power 3.1.9 software to perform statistical power analyses. Tumour burden (including number and/or size of lesions) was established as the primary outcome measure. Animal weights were monitored 3 times a week, and humane endpoints for euthanasia were established, including the assessment of the following parameters starting at 7 weeks of age: anemia, rectal bleeding, rectal prolapse development and low body condition score. Upon clinical presentation of any of these indicators by one of the animals in the cage (regardless of genotype), the whole cohort was sacrificed. The researchers involved in executing the procedures are certified in animal experimentation (FELASA C).

### CRC patient samples

Formalin-fixed paraffin-embedded (FFPE) CRC tissue specimens and normal colonic tissues were obtained from the FFPE tumour tissue biobank of the Centro Hospitalar Universitário de S. João (CHUSJ), Porto, Portugal. The samples were collected between January 2002 and December 2010, and their use for research purposes was approved by the CHUSJ Ethics Committee. Patient-derived organoids were established in collaboration with the Tissue Biobank of IPO Porto, Porto, Portugal, and approved by the IPO Ethics Committee (CES IPO: 81/022). A cohort of treatment-naïve adult patients (over 50 years old) was recruited between May 2022 and August 2023. Written informed consent was obtained from all patients prior to enrolment.

### Histochemical procedures

FFPE mouse tissue sections with 5μm (ISH) or 3μm (other procedures) thickness were deparaffinized and rehydrated following standard protocols. For morphological analysis, slides were stained with 7211 Haematoxylin for 1min, differentiated in 1% ammoniacal water for 30sec and counterstained using Eosin solution for 3min. For the AB-PAS staining, tissues were first stained with 1% AB (pH 2.5) for 45min, followed by treatment with 0.5% periodic acid for 10min, and then incubated with Schiff’s reagent for 15min. Nuclei were counterstained with Modified Mayer’s Haematoxylin for 2min. For IHC, heat-induced epitope retrieval was performed in a steamer (Black & Decker) for 40min with 10mM citrate buffer solution (pH=6) or 10mM Tris-EDTA solution (pH 8.0) unmasking solution, followed by 20min cooling at room temperature. Endogenous peroxidase activity was blocked with 3% H_2_O_2_ aqueous solution for 10min. For murine tissue sections, antibody non-specific interactions were blocked for 30min with normal rabbit or swine serum (1:5) in Antibody Diluent OP Quanto (Thermo Fisher Scientific), depending on the species of the secondary antibody. Slides were incubated with primary antibody overnight at 4°C. Tissues were then incubated with the corresponding biotinylated secondary antibody for 30min, followed by signal amplification with an avidin/biotin detection system (Vectastain ABC kit, Vector Laboratories) for 30min. For human tissues, the REAL EnVision Detection System, Peroxidase/DAB (Dako) was used after primary antibody incubation. Diaminobenzidine (DAB) was used as chromogenic substrate, with an incubation time of 5min. Nuclear contrast was performed with Modified Mayer’s Hematoxylin for 1min. All slides were dehydrated, clarified and permanently mounted with a xylene-based mounting medium.

### RNA *in situ* hybridization

The single-plex RNAscope 2.5 HD-RED ISH assay (Advanced Cell Diagnostics, Bio-Techne) was performed in FFPE tissue sections according to the manufacturer’s instructions, with the following modifications: (i) after deparaffinization and rehydration, sections were subjected to the mild pre-treatment protocol; (ii) AMP5 incubation was performed for 40 and 45min for human *LGR5* (Hs-LGR5, ref. 311021) and mouse *Lgr5* (Mm-Lgr5, ref. 312171), respectively, and for 1h for mouse *Mex3a* (Mm-Mex3a-E2-CDS, ref. 318551); and (iii) sections were counterstained with fast green stain solution (Thermo Fisher Scientific) for 1min. Slides were dried for 15min at 60°C, clarified and permanently mounted with Vectamount (Vector Laboratories). Incubations at 40°C were performed in a HybEZ hybridization oven (Advanced Cell Diagnostics, Bio-Techne).

### Conditioned media production for organoid cultures

Conditioned media (CM) for organoid cultures was produced in-house using a HEK293T cell line stably transfected with a HA-mouse RSPO1-Fc vector (a kind gift from Calvin Kuo, Stanford University, USA), a HEK293T cell line stably transfected with a mouse NOG-Fc vector (a kind gift from Gijs van den Brink and Vanesa Muncan, Amsterdam University Medical Centers, the Netherlands), and a commercial murine L cell line stably transfected with a WNT3A expressing vector (ATCC CRL2647). All cell lines were routinely cultured in T75 flasks with DMEM (Thermo Fisher Scientific) + 10% (v/v) FBS (Biowest) with appropriate selection antibiotics. For CM production, these were split (1:5) into 4 T75 flasks with complete DMEM without selection antibiotics. When reaching confluence, cells from 2 T75 flasks were combined into one T175 culture flask with Advanced DMEM/F12, 10% (v/v) FBS, 10mM HEPES and 1x GlutaMAX (both Thermo Fisher Scientific). After 7 days, CM was harvested, filtered with a 0.22μm PES filter and stored at −20°C until use.

### Mouse-derived CRC tumoroids culture

Focal lesions were isolated from the distal mouse colon and cut into small pieces of around 5mm^2^. The epithelial tissue was enzymatically separated from the underlying mesenchymal compartment by incubation with ACCUMAX cell detachment solution plus 10µM Y-27632 ROCK inhibitor (both from StemCell Technologies) for 30min at 37°C with agitation. The cell suspension was centrifuged and single cells embedded in Matrigel growth factor reduced basement membrane matrix (Corning). The cell/matrix suspension was plated in each well of a pre-warmed 24-well plate and incubated for 30min at 37°C, 5% CO_2_. After complete polymerization, 500µL of murine tumoroid medium was added to each well, composed of Advanced DMEM/F12, 10mM HEPES, 1x GlutaMAX, 1x N-2 Supplement, 1x B-27 Supplement (all Thermo Fisher Scientific), 100μg/mL primocin (InvivoGen), and 10% (v/v) NOG-CM, supplemented with 10µM Y-27632 only during establishment and passaging. Culture media was replaced every 2 days and tumoroids passaged every week. Assays were conducted after the culture had stabilized (at least three passages). Tumoroids size, morphology, and molecular marker expression were assessed after 7 days of culture. For PPARγ pathway activation, tumoroids were treated with 50µM Rosiglitazone (ROSI, Tocris Bioscience, Bio-Techne) or vehicle (DMSO) for 5 days.

### Patient-derived CRC tumoroids culture

For human patient-derived CRC tumoroid cultures, tumour and matched normal-like tissues obtained from patients undergoing surgery with curative intent were enzymatically processed with a Collagenase Type XI (Merck), DNAse I and Dispase (both from StemCell Technologies) cocktail solution, plus 10µM Y-27632, for 30min at 37°C under agitation. The cell suspension was centrifuged and single cells embedded in Matrigel as indicated above. Normal colon organoids were maintained in human colon organoid media composed of Advanced DMEM/F12, 10mM HEPES, 1x GlutaMAX, 1x N-2 Supplement, 1x B-27 Supplement, 1.25mM N-acetylcysteine (StemCell Technologies), 10nM gastrin I (Merck), 5μM SB202190 (Merck), 500nM A83-01 (Tocris Bioscience, Bio-Techne), 50% (v/v) WNT3A-CM, 10% (v/v) NOG-CM, 10% (v/v) RSPO1-CM, 50ng/mL EGF (Peprotech, Thermo Fisher Scientific), 100μg/mL primocin, 1µM CHIR99021 (StemCell Technologies) and 10µM Y-27632 (only during establishment and passaging). CRC-derived tumoroids were kept in human CRC tumoroid media, which is similar to human colon organoid media but without the addition of WNT3A-CM and CHIR99021. Culture media was replaced every 2 days and tumoroids passaged every two weeks. Assays were conducted after the culture had stabilized (at least three passages). Tumoroids size, morphology, and molecular marker expression were assessed after 14 days of culture. To address chemosensitivity, tumoroids were plated in 96-well culture plates and treated with 5-Fluorouracil (5-FU; Merck) plus Oxaliplatin (MedChemExpress) in a 25:1 ratio equivalent to the combined IC50 dose (282.5µM 5-FU and 11.3µM Oxaliplatin) or vehicle-treated for 72h. Cell viability was measured using the CellTiter-Glo 3D Cell Viability assay (Promega) following the manufacturer’s instructions.

### Establishment of CRISPR-Cas9-mediated MEX3A KO tumoroid lines

A single-guide (sg)RNA targeting the MEX3A coding region (https://genome.ucsc.edu/cgi-bin/hgTracks) was cloned into a SpCas9-EGFP vector (PX458; Addgene #48138) using a previously described protocol^53^ The patient-derived CRC tumoroid line 005T was dissociated and plated in collagen type I (ibidi) coated plates. Adherent cells were transfected with 4μg of the PX458-sgMEX3A expressing vector using Lipofectamine 2000 reagent (Thermo Fisher Scientific). Cells were enzymatically removed from the collagen 48h after transient transfection, single EGFP+ cells collected by fluorescence-activated cell sorting (FACS) using a BD FACS Aria II sorter (BD Biosciences) and plated in collagen type I coated 96-well plates to establish monoclonal populations of putatively edited tumour cells. Downstream analysis was performed after reestablishment of the culture in Matrigel. KO efficiency of different clones was evaluated by Western blot analysis of MEX3A protein expression and by DNA Sanger sequencing.

### Protein extraction and Western Blot

Cells were lysed on ice for 30min with lysis buffer containing 20mM Tris-HCl (pH=7.5), 150mM NaCl, 2mM EDTA, 0.1% (v/v) sodium deoxycholate and 0.1% (v/v) sodium dodecyl sulfate supplemented with 1x Complete protease inhibitor cocktail (Roche Applied Science), 20mM NaF, 1mM PMSF and 1mM Na_3_VO_4_. Lysates were centrifuged for 15min at 17000x*g* at 4°C and the supernatant recovered. Protein concentration was determined with BCA Protein Assay kit (Thermo Fisher Scientific). 20-50μg of each protein extract were run in a 10% SDS-PAGE gel, transferred to a nitrocellulose membrane and incubated overnight with the desired antibodies after which signals were revealed with ECL detection kit (GE Healthcare Life Sciences). Actin or GAPDH levels were used to normalize protein expression, and quantification was performed using Fiji software.^54^

### Site-directed mutagenesis and HyperTRIBE constructs

A previously published MEX3A expression vector^25^ was used together with a *Drosophila melanogaster* ADARcd expression vector (a kind gift from Jean-Pierre Rouault, Centre de Recherche en Cancérologie de Lyon, France). An inactive MEX3A RNA-binding mutant (MEX3AΔKH) was generated by introducing a double amino acid change in both KH domains: R150D and Q151D in KH1, P241D and K242D in KH2, previously predicted to impair nucleic acid binding without causing significant structural changes or compromising domain stability.^55^ The hyperactive version of *Drosophila’s* ADARcd was generated by introducing E458Q, the human ADAR2 E488Q matching mutation, previously described to increase human’s ADAR2cd editing efficiency and reduce sequence recognition bias.^56, 57^ All mutations were introduced with the Q5 Site-Directed Mutagenesis Kit (New England Biolabs) using oligonucleotides containing the desired mutations. The ADARcd, MEX3A and MEX3AΔKH coding sequences were amplified with overlapping primers, allowing assembly of adjacent fragments. Amplified fragments were cloned into the pCW57-RFP-P2A-MCS lentiviral vector (Addgene #78933) after *BamH*I/*Pst*I digestion, using the NEBuilder High-Fidelity DNA Assembly Cloning Kit (New England Biolabs), generating MEX3A-ADARcd and MEX3AΔKH-ADARcd HyperTRIBE vectors. Sanger sequencing was performed to confirm proper cloning using the BigDye Terminator v3.1 Cycle Sequencing Kit (Thermo Fisher Scientific). Sequencing products were purified with Sephadex G-50 Fine DNA Grade (GE Healthcare) before analysis.

### Lentiviral particles production and SW480 cell transduction

Human embryonic kidney (HEK)293T (ATCC CRL-3216) cell line was used for production of replication-incompetent lentiviral particles to deliver the HyperTRIBE vectors. Briefly, 2μg psPAX2 packaging vector (Addgene #12260), 2μg pCMV-VSV-G envelope vector (Addgene #14888) and 4μg MEX3A-ADARcd or MEX3AΔKH-ADARcd vectors were diluted in Opti-MEM (Thermo Fisher Scientific) and transiently transfected into HEK293T cells using Lipofectamine 2000 reagent. Viral particle-containing supernatants were obtained 48h later, filtered through a 0.45μm PES filter, and precipitated with PEG-it Virus precipitation kit (System Biosciences). The human colorectal carcinoma cell line SW480 (ATCC CCL-228) was plated and left for 24h in DMEM + 10% (v/v) FBS to adhere. The media was then replaced by fresh media containing diluted viral supernatant plus 15μg/mL polybrene (Merck). After incubation for 96h, the media was removed and selection of transduced cells initiated with 10μg/mL puromycin (InvivoGen). SW480 HyperTRIBE stable cell lines were plated for system induction with 1µg/mL doxycycline. After 48h of treatment, TurboRFP+ cells were collected by FACS, along with non-induced cells, pelleted and stored at −80°C for downstream analysis.

### RNA extraction and reverse transcription quantitative PCR

For the HyperTRIBE assay, RNA extraction was performed using the TRIzol Plus Total Transcriptome isolation protocol, combining the TRI reagent (Merck) and PureLink RNA Mini Kit (Thermo Fisher Scientific). For reverse transcription quantitative PCR, RNA was extracted using only the TRI Reagent according to the manufacturer’s instructions. RNA was quantified using a NanoDrop 1000 spectrophotometer (Thermo Fisher Scientific) and 2µg of total RNA reverse-transcribed using the NZY Reverse Transcriptase (NZYTech). Analysis of mRNA expression was performed by quantitative real-time PCR in an ABI prism 7500 system with PowerUp SYBR Green Master Mix (Thermo Fisher Scientific) and specific primer pairs. Each sample was quantified in triplicate and specificity confirmed by dissociation analysis. Gene expression was calculated through the relative standard curve method, with *18S* rRNA levels used for target gene abundance normalization.

### RNA-sequencing

SW480 HyperTRIBE total RNA quantification and quality control were assessed using a 2100 Bioanalyzer instrument (Agilent Technologies). Only samples with an RNA integrity number (RIN) above 9 were considered for further analysis. Briefly, RNA samples were processed for rRNA removal using the Ribo-Zero kit (Illumina). Then, mRNA was randomly fragmented by adding fragmentation buffer and cDNA synthesized using random hexamers, after which a custom second-strand synthesis step was performed. This was followed by purification with AMPure XP beads, terminal repair, polyadenylation, sequencing adapter ligation, size selection and degradation of second-strand U-contained cDNA by the USER enzyme. After the final PCR enrichment, a strand-specific cDNA library was generated using the NEBNext Ultra II Directional RNA Library Prep kit for Illumina (New England Biolabs). These procedures were performed by Novogene (UK). Two biological replicates of each condition were used for stranded sequencing with 150bp paired-end reads on the NovaSeq platform (Illumina). An average of 86 million reads were generated per sample.

### Bioinformatic analysis of RNA-sequencing data

Trimming of the reads was first carried out using Trimmomatic version 0.39, with the following non-default settings recommended for the HyperTRIBE pipeline (https://github.com/rosbashlab/HyperTRIBE): Head-Crop: 6; Number of leading bases for trimming: 25; Number of trailing bases for trimming: 25; Average quality of 3 consecutive bases: 25; Minimum length 19; Adapters trimming (using TruSeq3 adapters): TruSeq3-PE.fa:2:30:10:2:keepBothReads. The human reference genome, build GRCh38 (patch release 12), and the curated gene structure were retrieved from the UCSC Genome Browser. The alignment of the paired-end RNA-seq reads to the human genome assembly was performed using STAR aligner (v2.7.8a).^58^ Besides default options, the following were applied specifically for HyperTRIBE: outFilterMismatchNoverLmax: 0.07; outFilterMatchNmin: 16; outFilterMultimapNmax: 1. The aligned reads were outputted in BAM format sorted by coordinates. Aligned reads were further filtered in samtools (v1.12), considering only alignments with a mapping Phred quality score above 10. Duplicate marking of reads was carried out in MarkDuplicates (Picard) tool using the “REMOVE_DUPLICATES” option. Next, we followed the GATK workflow^59^ for variant calling in RNA-seq to identify mutations in the aligned reads. Prior to the variant calling step, as per the GATK recommendations, a base quality recalibration step was carried out in order to detect and fix systematic errors on the quality scores. We then restricted the analysis to mutations within annotated RNA transcripts, retrieved from the NCBI RNA reference sequence (RefSeq) collection, calling out A-to-G mutations in transcripts encoded by the forward strand and T-to-C mutations in transcripts encoded by the reverse strand. For unambiguous identification of RNA-edited sites above background, we considered an A frequency > 80%, a G frequency of 0% in the control RNA, and a G frequency > 0% in the experimental RNA. The filtered sets of RNA editing events from RNA-seq libraries of the same experimental condition were combined and the number of reads containing the reference (A or T) and alternative (G or C) alleles from each library at each site were counted. We applied a beta-binomial distribution to model RNA edit frequencies,^60^ comparing the MEX3A-ADARcd induced condition against both the MEX3AΔKH-ADARcd induced and the non-induced control samplesThe *P* values from all sites were adjusted to control for false discovery rate (FDR) using a Benjamin-Hochberg correction. Significant sites were determined by filtering for FDR-adjusted<0.001, an expression level of at least 1 FPKM and a minimum of 20 reads counts in each MEX3A-ADARcd replicate. Edited sites were required to be present in both replicates. High confident target genes were retained considering no editing in controls (G or C = 0) and a differential edit frequency of at least 25%. For *de novo* motif discovery, we extracted the sequences extending 500bp from both sides of each edit site in the 3’UTR regions of the previously identified transcripts and considered these windows as the target sequence pool for the HOMER software. Overlapping sequences were merged into a single sequence and background sequences of similar length were randomly selected from 3’UTRs in the genome that did not overlap with the target sequence. The search was limited to enriched motifs of 6, 7, or 8 nucleotides in length, and a regional oligomer autonormalization of up to 3 nucleotides in length. For differential gene expression analysis, the aligned reads previously mapped were counted using HTSeq-Count.^61^ Differential expression analysis between paired tests were identified by the DESeq2 package, which applies a negative binomial distribution test to model gene counts and test for differential expression. Only genes with a *P* value threshold adjusted for multiple testing (Benjamini-Hochberg method) <0.01 were considered.

### Statistical analyses

At least three biological replicates were done for each experiment, with three replicates per assay. Statistical analysis of data was performed using the GraphPad Prism software and differences between groups considered statistically significant at a *P* < 0.05. For statistical analysis of differences between tumour burden in mice the Chi-Square test was applied, while differences in tumoroid morphology were accessed through Fisher’s exact test. Differences for tumour area, tumoroid diameter, wall thickness, and response to chemotherapy were determined using two-tailed unpaired *t*-test or Mann Whitney test for data not following normal distribution. Differences in mRNA expression and number of editing events in HyperTRIBE were evaluated with one-way ANOVA. For human CRC tissues, results are expressed as a percentage or mean ± standard deviation (SD). For analysis of the relationship between patient’s age and biomarker expression, we used two-tailed unpaired *t*-test. Pearson’s Chi-Square test was used in the statistical analysis of parameters where all classes had an expected count above 10, otherwise, Fisher’s exact test was implemented.

## ACKNOWLEDGMENTS

The authors thank Sérgia Velho (i3S, Porto, Portugal) for sharing the animal strains. The authors thank Sofia Lamas and all the technical staff at the i3S Animal Facility for assistance with animal experiments. The authors acknowledge the support of the i3S Scientific Platform Histology and Electron Microscopy (HEMS), member of the national infrastructure PPBI - Portuguese Platform of Bioimaging (PPBI-POCI-01-0145-FEDER-022122), as well as the Bioinformatics and Genomics i3S Scientific Platforms. We thank Jean-Pierre Rouault (Centre de Recherche en Cancérologie de Lyon, Lyon, France) for providing the ADARcd coding sequence expression vector. We also thank Katharine Abruzzi (Howard Hughes Medical Institute, Brandeis University, Waltham, MA, USA) and Jeetayu Biswas (Memorial Sloan Kettering Cancer Center, New York, NY, USA) for their helpful insights regarding the HyperTRIBE technique.

## CREDIT AUTHORSHIP CONTRIBUTIONS

**Ana R. Silva**, M.Sc. (Methodology: Supporting; Investigation: Lead; Validation: Lead; Writing – original draft: Supporting; Writing - Review & Editing: Supporting; Visualization: Supporting); **Alexandre Coelho**, M.Sc. (Methodology: Supporting; Investigation: Supporting; Validation: Supporting; Writing - Review & Editing: Supporting); **Vanessa Machado**, Ph.D. (Methodology: Supporting; Investigation: Supporting; Validation: Supporting; Writing - Review & Editing: Supporting); **Morgana Russel**, M.Sc. (Investigation: Supporting); **Dalila Mexieiro**, M.Sc. (Investigation: Supporting); **Ana L. Amaral**, M.Sc. (Resources: Supporting; Writing - Review & Editing: Supporting); **Bruno Cavadas**, Ph.D. (Software: Lead; Formal analysis: Lead; Writing - Review & Editing: Supporting); **Carina Carvalho-Maia** (Resources: Supporting); **Davide Gigliano** (Resources: Supporting); **Carmen Jerónimo**, Ph.D. (Resources: Lead; Writing - Review & Editing: Supporting); **Raquel Almeida**, Ph.D. (Methodology: Supporting; Writing - Review & Editing: Supporting; Supervision: Supporting; Funding acquisition: Supporting); **Bruno Pereira**, Ph.D. (Conceptualization: Lead; Methodology: Lead; Investigation: Supporting; Validation: Supporting; Writing - original draft: Lead; Writing - Review & Editing: Lead; Visualization: Lead; Supervision: Lead; Project administration: Lead; Funding acquisition: Lead).

## FUNDING

This work was supported by Portuguese funds through FCT (Fundação para a Ciência e a Tecnologia, I.P.) within the framework of the diffNET research project (https://doi.org/10.54499/PTDC/BIA-CEL/0456/2021). Bruno Cavadas and Bruno Pereira acknowledge FCT for financial support through the CEEC program (https://doi.org/10.54499/CEECINST/00123/2021/CP1772/CT0001 and https://doi.org/10.54499/CEECINST/00131/2021/CP2805/CT0002, respectively). Rita Silva and Alexandre Coelho acknowledge FCT for financial support through PhD fellowships (https://doi.org/10.54499/2021.06919.BD and 2022.10128.BD, respectively).

## COMPETING INTERESTS

Bruno Pereira received grant funding from Merck KGaA (Darmstadt, Germany). The remaining Authors have no conflicts of interest to disclose.

## DATA AVAILABILITY STATEMENT

The Authors declare that all supporting data are available within the paper and its Supplementary Material, or from the corresponding Author on reasonable request. Raw RNA-seq data will become available in the Gene Expression Omnibus (GEO) database upon publication.

## FIGURES

**Supplementary Figure S1.**
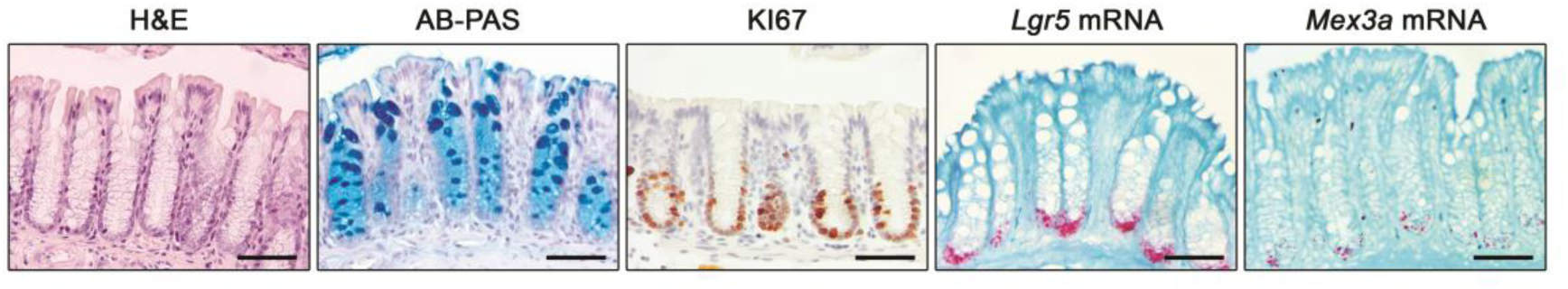
*Mex3a* transcripts accumulate at the bottom of the crypts in wild-type mice. Representative histology of colon from wild-type adult mice (H&E and AB-PAS staining, KI67 IHC, *Lgr5* and *Mex3a* mRNA ISH). Scale bars, 50µm.

**Supplementary Figure S2.**
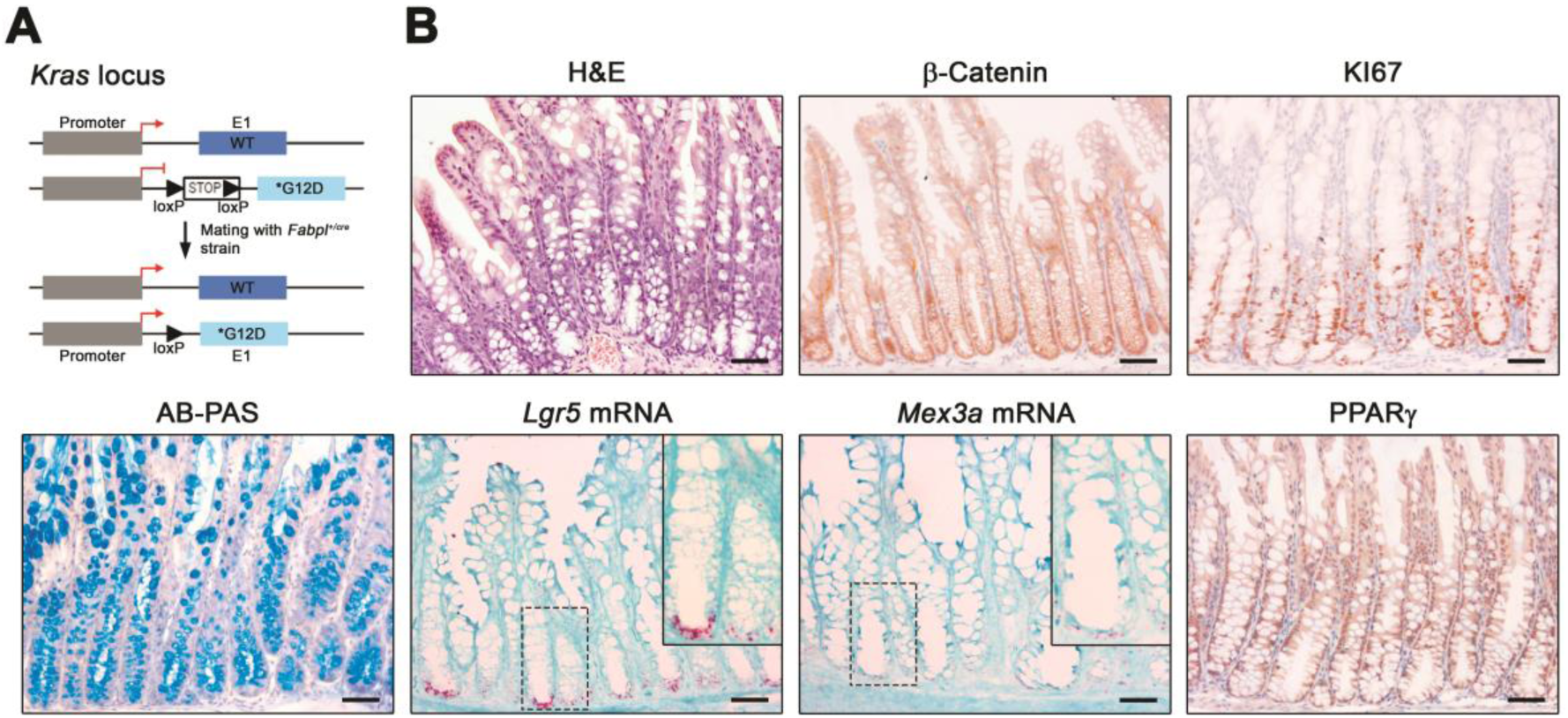
Colon of *Kras^+/G12D^*;*Fabpl^+/Cre^* mice presents widespread hyperplasia. **A,** Schematic representation of the genetically modified *Kras* locus for conditional expression of the constitutively active *Kras^G12D^* mutant upon Cre-mediated recombination under the control of the *Fabpl* gene promoter. **B,** Representative histopathology of *Kras^+/G12D^*;*Fabpl^+/Cre^* colonic mucosa (H&E and AB-PAS staining; β-Catenin, KI67 and PPARγ IHC; *Lgr5* and *Mex3a* mRNA ISH). Inserts depict high magnification of the boxed areas. Scale bars, 100µm.

**Supplementary Figure S3.**
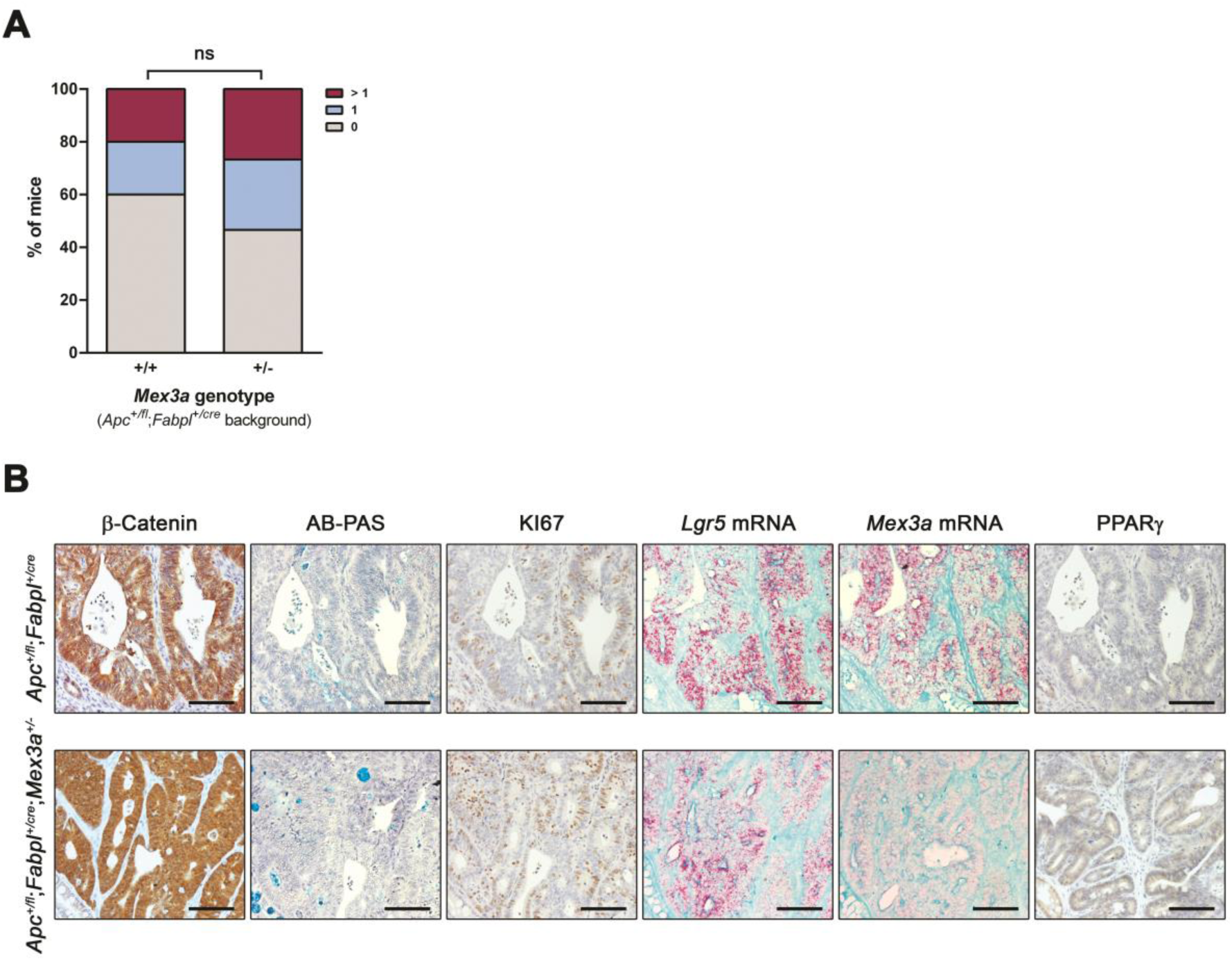
*Mex3a* levels do not impact tumour burden in the colon of *Apc^+/fl^;Fabpl^+/cre^*mice. **A,** Quantification of the number of tumours in the colon of *Apc^+/fl^*;*Fabpl^+/cre^* and *Apc^+/fl^*;*Fabpl^+/cre^*;*Mex3a^+/−^* mouse models (n = 15 for each genotype). ns (not significant) *P* = 0.7650, Pearson Chi-square test. **B,** Representative histopathology of *Apc^+/fl^*;*Fabpl^+/cre^*and *Apc^+/fl^*;*Fabpl^+/cre^*;*Mex3a^+/−^*colonic mucosa lesions (AB-PAS staining, β-Catenin, KI67 and PPARγ IHC, *Lgr5* and *Mex3a* mRNA ISH). Scale bars, 50µm.

**Supplementary Figure S4.**
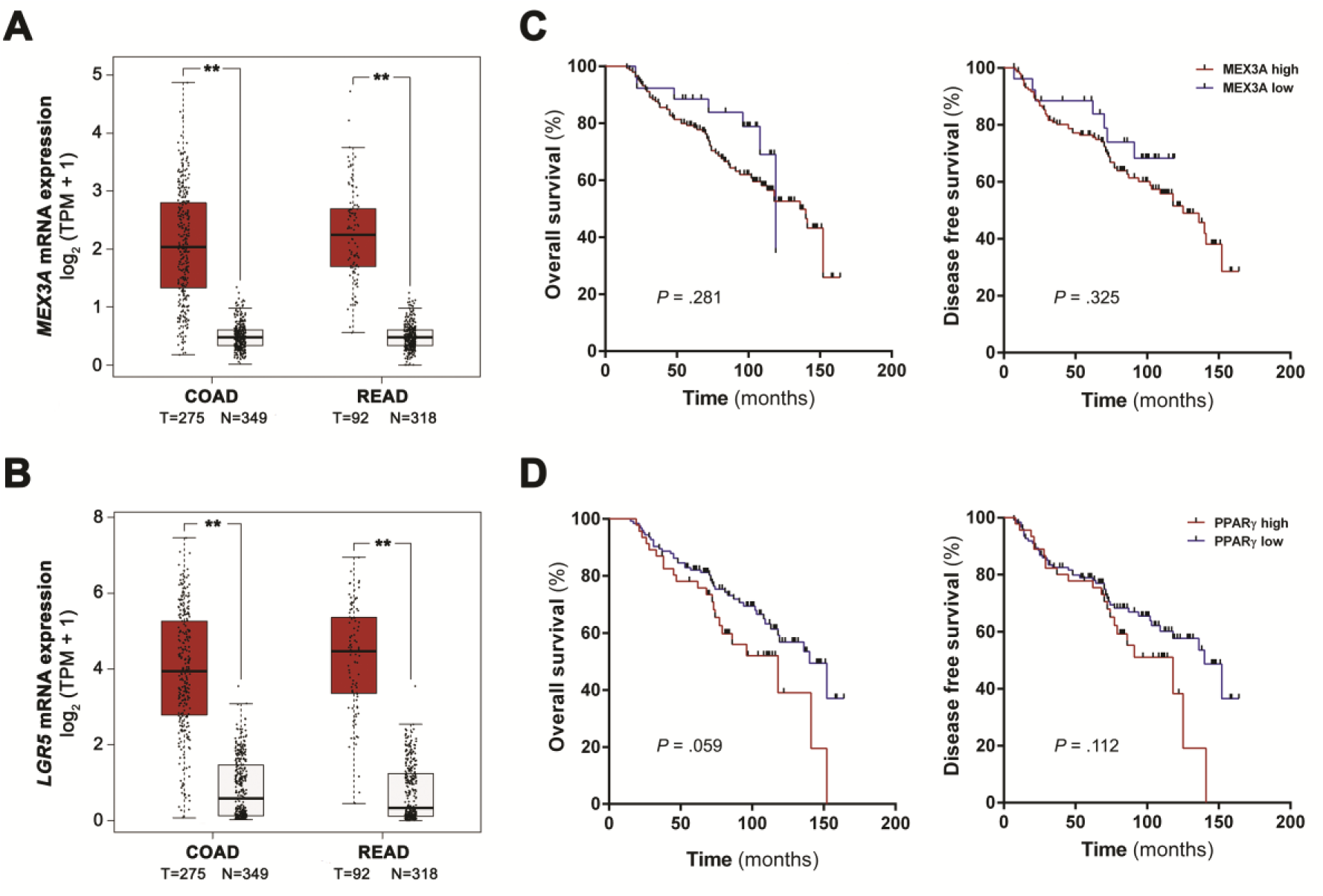
ISC markers *MEX3A* and *LGR5* are overexpressed in CRC human tissues. **A** and **B**, Analysis of *MEX3A* **(A)** and *LGR5* **(B)** mRNA expression levels from TCGA and the non-disease state GTEx portal (expression in log_2_ scale). COAD - Colon adenocarcinoma; READ - Rectal Adenocarcinoma; T - tumour; N – normal. \*\**P* < 0.01, one-way ANOVA test. **C** and **D**, Kaplan-Meier curves of overall and disease-free survival according to MEX3A **(C)** and PPARγ **(D)** expression levels in a cohort of CRC stage II patients. p-values depicted in the respective plot, log-rank (Mantel-Cox) test.

**Supplementary Figure S5.**
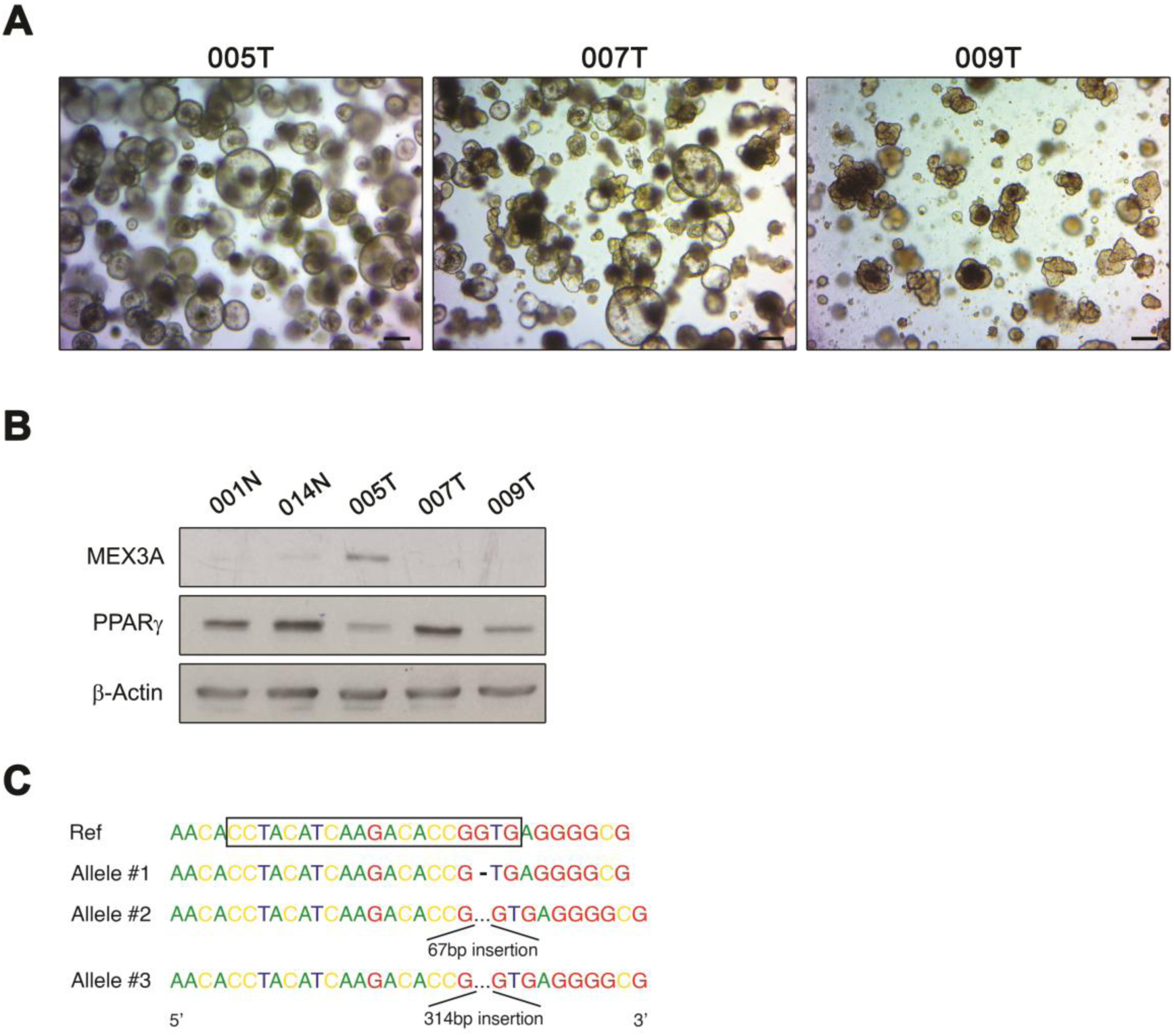
PDCT and normal colon-derived organoid lines present inversely correlated MEX3A and PPARγ protein expression profiles. **A,** Representative phase-contrast microscopy images of three PDCT lines after 14 days of culture. Scale bars, 200µm. **B,** Representative western blots of PDCTs and normal colon-derived organoid lines for MEX3A and PPARγ expression. **C,** Schematic representation of the *MEX3A* alleles in the 005T_AE6 tumoroid line identified upon cloning on pGEM®-T Easy vector and Sanger sequencing. Boxed area highlights the region targeted by the sgRNA. No alterations were detected in the 005T line.

**Supplementary Figure S6.**
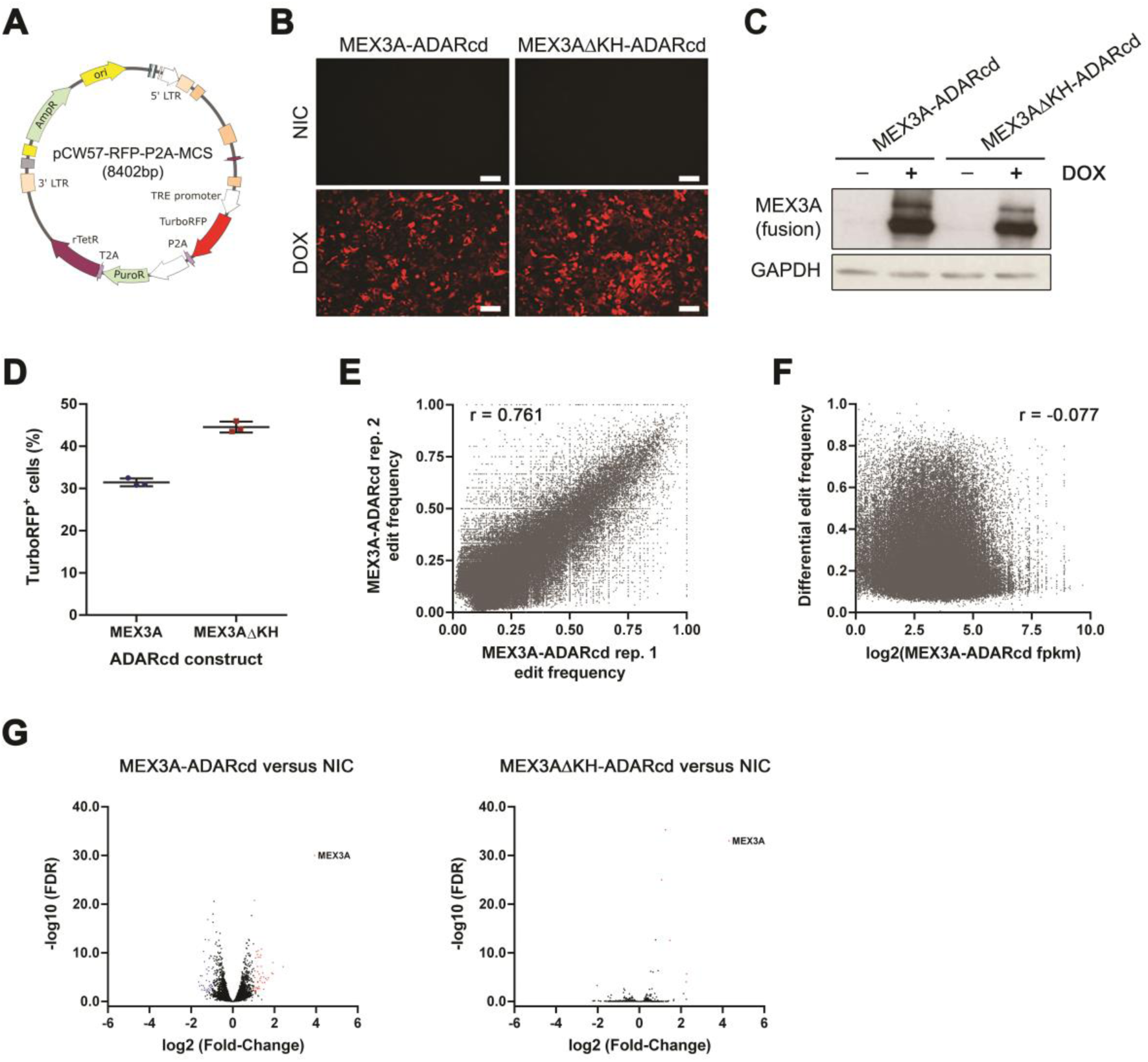
HyperTRIBE system implementation and quality control in SW480 CRC cells. **A,** Graphic representation of the vector used to express MEX3A- ADARcd and MEX3AΔKH-ADARcd fusion proteins. **B,** Representative images of MEX3A-ADARcd and MEX3AΔKH-ADARcd stably transduced cell lines after 48h of HyperTRIBE system induction with 1µg/mL DOX and respective NIC control. **C,** Representative western blot of MEX3A fusion proteins in HyperTRIBE system induction experiments in the SW480 cell line. MEX3A-ADARcd fusion proteins have a higher molecular weight (predicted around 100kDa) than the endogenous MEX3A protein (65kDa). **D,** Average proportion of TurboRFP^+^ cells after 48h of HyperTRIBE system induction (3 independent induction experiments). **E,** Linear correlation analysis between the two independent replicates for editing frequency. r = 0.761. **F,** Linear correlation analysis between editing frequency and transcript expression level (fpkm). r = −0.077. **G,** Volcano plots of the differential expressed genes; blue and red dots represent genes with down or upregulated expression (respectively) in MEX3A-ADARcd or MEX3AΔKH- ADARcd versus NIC control.

**Supplementary Table 1.**
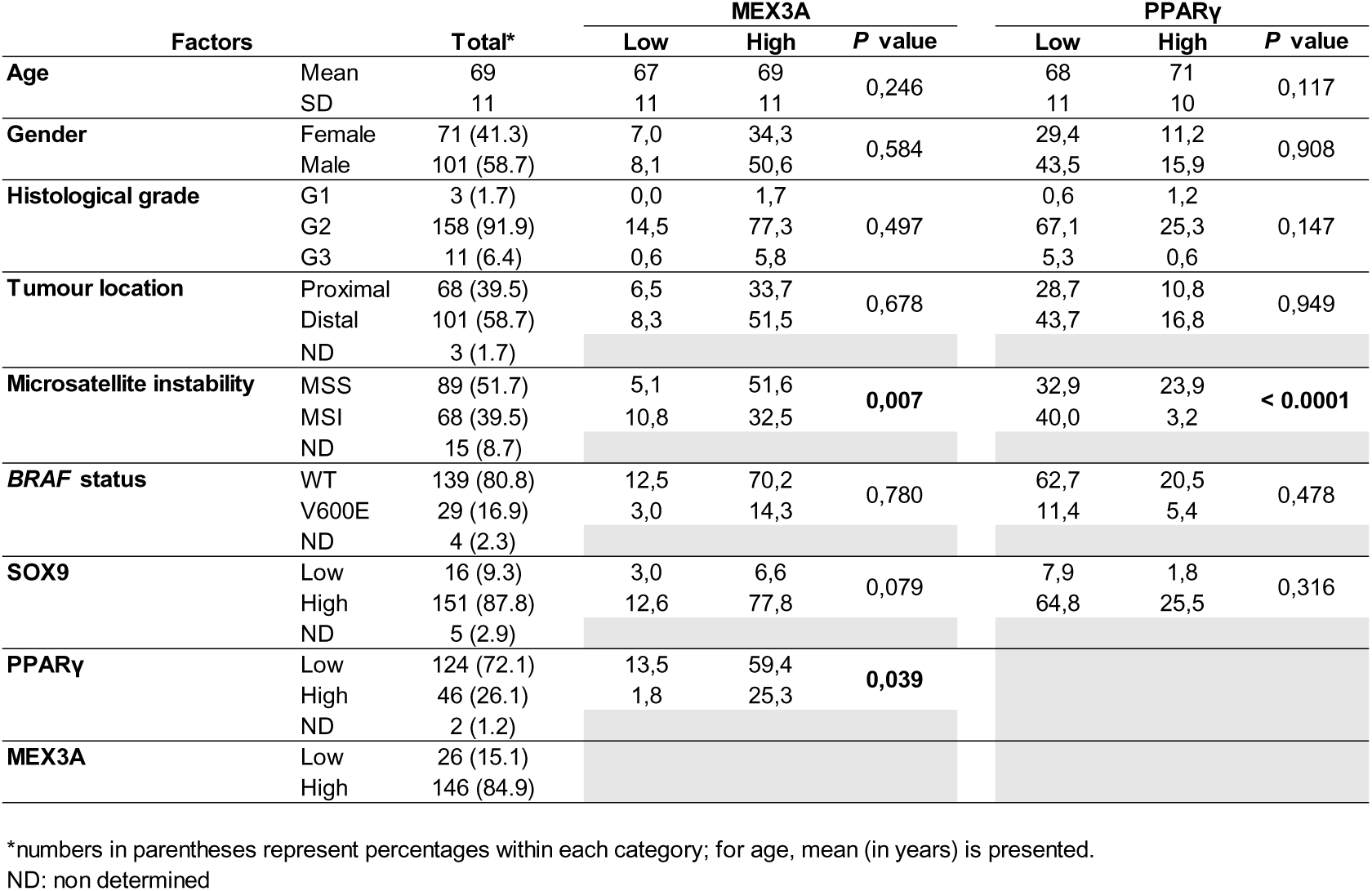
Summary of clinicopathological and molecular associations with MEX3A and PPARγ expression levels in colon cancer. Results are expressed as a percentage or mean plus standard deviation (SD).

## Notes

### Competing Interest Statement

Bruno Pereira has received grant funding from Merck KGaA, Darmstadt, Germany. The authors have no additional financial interests.

